# Identification of a Novel Hypoxia-induced Inflammatory Cell Death Pathway

**DOI:** 10.1101/2023.08.05.552118

**Authors:** Abhishek Bhardwaj, Maria Antonelli, Beatrix Ueberheide, Benjamin G. Neel

**Affiliations:** Laura and Isaac Perlmutter Cancer Center, New York University Grossman School of Medicine, NYU Langone Health, New York, NY, USA; Proteomics Laboratory, Division of Advanced Research Technologies, New York University Grossman School of Medicine, New York, NY, USA; Department of Biochemistry and Molecular Pharmacology, New York University Grossman School of Medicine, New York, NY, USA; Department of Neurology, New York University Grossman School of Medicine, New York, NY, USA

**Keywords:** RNF213, PTP1B, CYLD, LUBAC, NF-κB, Hypoxia, Inflammation, ER-stress, Pyroptosis, MMD

## Abstract

Hypoxic cancer cells resist many anti-neoplastic therapies and can seed recurrence. We found previously that PTP1B deficiency promotes HER2+ breast cancer cell death in hypoxia by activating RNF213, an ∼600kDa protein containing AAA-ATPase domains and two ubiquitin ligase domains (RING and RZ) that also is implicated in Moyamoya disease (MMD), lipotoxicity, and innate immunity. Here we report that PTP1B and ABL1/2 reciprocally control RNF213 phosphorylation on tyrosine-1275. This phosphorylation promotes RNF213 oligomerization and RZ domain activation. The RZ domain ubiquitylates CYLD/SPATA2, and together with the LUBAC complex, induces their degradation. Decreased CYLD/SPATA2 causes NF-κB activation, which together with hypoxia-induced ER-stress triggers GDSMD-dependent pyroptosis. Mutagenesis experiments show that the RING domain negatively regulates the RZ domain. *CYLD*-deleted HER2+ cell-derived xenografts phenocopy the effects of PTP1B deficiency, and reconstituting *RNF213* knockout lines with RNF213 mutants shows that the RZ domain mediates PTP1B-dependent tumor cell death. Our results identify a novel, potentially targetable PTP1B/RNF213/CYCLD/SPATA pathway critical for controlling inflammatory cell death in hypoxic tumors that could be exploited to target hypoxic tumor cells, potentially turning “cold” tumors “hot”. Our findings also reveal new insights into RNF213 regulation, and have potentially important implications for the pathogenesis of MMD, atherosclerosis, and inflammatory and auto-immune disorders.

## Main

Most tumors have hypoxic or anoxic areas^1^. In response, cancer cells adapt to limited oxygen in the tumor microenviroment (TME) by successively activating several signaling pathways. In normoxia, the transcription factors HIF1α and HIF2α (hereafter “HIF”) are hydroxylated by oxygen-dependent prolyl hydroxylases (PHD 1-3), bound by the Van Hippel Lindau tumor suppressor gene product (VHL), and targeted for ubiquitylation and proteosomal degradation^2^. PHD enzymes are inhibited in low oxygen, resulting in HIF stabilization and induction of hypoxia response genes. More severe hypoxia (≤1% O_2_) triggers the AMP-dependent kinase (AMPK) and endoplasmic reticulum (ER) stress pathways, which evoke distinct adaptive responses^3,4^. The activity of NF-κB, a key transcription factor controlling inflammation and innate immunity, also increases in hypoxia via (an) incompletely defined mechanism(s) that include(s) decreased PHD regulation of NF-κB pathway signaling components^5^. Conversely, NF-κB is required for basal *HIF1alpha* expression, and thus directly and indirectly regulates the transcriptome of hypoxic cells^6^. The complex interplay between HIF-induced, metabolic (AMPK), integrated stress response (ER stress), and inflammatory/innate immune (NF-κB)^7,8^ pathways helps determine cancer cell sensitivity to conventional (chemoradiation) and targeted therapy^3^, as well as whether tumor cell death evoked by antineoplastic therapies is “sterile” or immunogenic^9–11^.

The protein-tyrosine phosphatase PTP1B (encoded by *PTPN1*) resides on the cytosolic surface of the endoplasmic reticulum (ER), where it dephosphorylates incoming receptor tyrosine kinases (RTKs) and cytokine receptors^12,13^. Best known as a negative regulator of insulin and leptin signaling^14–17^, PTP1B is also required for *Neu* (encoding rodent HER2)-induced breast cancer in mice^18,19^. Moreover, *PTPN1* is amplified (∼5%) and overexpressed (∼72%) in human cancer^20,21^. Previously, we reported that PTP1B deficiency or inhibition increases the death of HER2+ breast cancer cells maintained at ≤1% O_2_ without altering the HIF, ER stress, or AMPK pathways^22^. Instead, hypoxia hypersensitivity required RNF213, a large (∼600 kDa) protein with AAA-ATPase and E3 ligase domains. PTP1B deficiency/inhibition promoted an increase in RNF213-dependent global ubiquitylation, and RNF213 was tyrosine phosphorylated and appeared to be a PTP1B substrate. However, key details, including how PTP1B regulates RNF213 and the mechanism and type of RNF213-dependent cell death, remained unclear.

RNF213 features six AAA-ATPase modules, two of which are active, a RING E3 ligase domain, a recently discovered “RZ” E3 domain, and large unstructured regions^23–27^. In an AAA-ATPase-and ISG15-dependent manner, RNF213 can oligomerize into a hexameric, ∼3.6mDa complex^23,25^. RNF213 localizes on lipid droplets^28^ (also dependent on AAA-ATPase activity/ISG15) and is required for lysophosphatidic acid (LPA)-induced cell death *in vitro*^29^. Transfection studies implicate RNF213 in NF-κB signaling^27,29^, and recently, the RNF213 RZ domain was reported to ubiquitylate lipid A of Salmonella^26^. In concert, these findings suggest roles for RNF213 in lipoxicity and inflammation, both potentially relevant for cancer.

We found that ABL1/2 phosphorylate, whereas PTP1B dephosphorylates, RNF213 on tyrosine 1275 (Y1275) in breast cancer cells. Phosphorylation of this site promotes RNF213 oligomerization and activation of its E3 ligase activity. We identify the CYLD-SPATA2 complex, a major negative regulator of NF-κB activation, as an RNF213 substrate critical for hypoxia-induced cell death. The RZ domain is required for CYLD-SPATA2 ubiquitylation/degradation, while the RING domain restrains RZ activity. Wild type RNF213 requires the LUBAC complex for effective CYLD-SPATA2 degradation, but RNF213 RING mutants, including highly penetrant SNPs associated with MMD, bypass this requirement. Increased RNF213 activity, caused by PTP1B deficiency/inhibition or RNF213 RING mutations, drives CYLD/SPATA2 ubiquitylation and degradation and increases NF-κB activity, priming cells for pyroptosis via the NLRP3 inflammasome. Concomitantly, hypoxia-induced ER stress delivers the “second signal” triggering cancer cell death. Deletion of SPATA2 phenocopies the effects of *PTPN1* deletion on tumorigenesis by HER2+ breast cancer cells. Our results have implications for the mechanism of action and potential uses of dual inhibitors of PTP1B/TC-PTP (*PTPN2*),^30^ now in clinical trials for various cancers as well as for the pathogenesis of MMD and other steno-occlusive disorders. Overall, we identify a new strategy to target hypoxic cancer cells that resist anti-neoplastic therapies.

## Results

### Y1275 phosphorylation of RNF213 is regulated by PTP1B/ABL, which promotes RNF213 oligomerization

We have reported that HER2+ breast cancer cells with *PTPN1* knockdown or treated with a PTP1B inhibitor are hypersensitive to hypoxia^22^. By studying an expanded panel of breast cancer cell lines, we found that hypoxia hypersensitivity is not limited to the HER2+ subtype; treatment with the allosteric PTP1B inhibitor claramine also caused hypoxia hypersensitivity in several ER+ and triple negative (TNBC) breast cancer lines (Extended Data Fig. 1a). Hence, PTP1B is more general regulator of the hypoxia response, at least in breast cancer cells and possibly in other malignancies.

We also found previously that RNF213 is tyrosine phosphorylated in HER2+ breast cancer cells and interacts with PTP1B. “Substrate-trapping” mutants of PTP1B show increased RNF213 binding, and this interaction is competed by the active site PTP inhibitor sodium orthovanadate, which suggested that RNF213 is a PTP1B substrate^22^. However, RNF213 tyrosine phosphorylation is not increased in PTP1B-inhibited/deficient HER2+ breast cancer cells, indicating that only select sites are PTP1B targets. We generated an RNF213-TurboID fusion protein with the goal of identifying RNF213 binding partners by proximity ligation^31^ (see below), labelled interacting proteins with biotin, enriched these proteins on streptavidin beads, performed on-bead tryptic digestion, and analyzed the peptide mixture by mass spectrometry (MS). This analysis revealed evidence of RNF213 phosphorylation on residue Y1275 (Supplemetary Table 1, Extended Data Fig. 1b).

To ask whether this phosphorylation is relevant for RNF213/PTP1B interaction, we used CRISPR/Cas9 to generate *RNF213*-knockout (KO) BT474 cells and retroviral gene transduction to introduce *PTPN1* or non-targeting (Control) shRNAs. We transduced these cells with an lentivirus expressing hemaglutinin (HA) tagged-mouse *Ptpn1*^D/A^, which encodes the substrate-trapping mutant PTP1B^D181A^ (HA-PTP1B^D/A^). The resultant lines were transfected with expression vectors for 3x-FLAG-*RNF213* or 3x-FLAG-*RNF213^Y^*^127^*^5F^*. Immunoprecipitation (IP) of HA-PTP1B^D/A^ resulted in recovery of 3x-FLAG-RNF213. Much less 3x-FLAG-RNF213^Y1275F^ co-immunoprecipitated (Fig. 1a, right), even though it was expressed at comparable levels to wild-type RNF213 (Fig. 1a, left). Similar results were obtained from analogous experiments with AU565-derived cell lines (Extended Data Fig. 1c). Tyrosine phosphorylation of 3x-FLAG-RNF213^Y1275F^ was reduced compared with that of 3x-FLAG-RNF213, but to a lesser extent (∼40%) than the near total decrease in 3x-FLAG-RNF213^Y1275F^ recovery in HA-mPTP1B^D/A^ immunoprecipitates (Compare Fig. 1a and 1b and Extended Data Fig. 1c, 1d). These data suggest that additional tyrosine phosphorylation sites exist in RNF213 that are not detected by MS or that new sites are generated in RNF213^Y1275F^. Regardless, our results indicate that RNF213-Y1275 is a specific site for PTP1B-catalyzed dephosphorylation.

**Figure 1.**
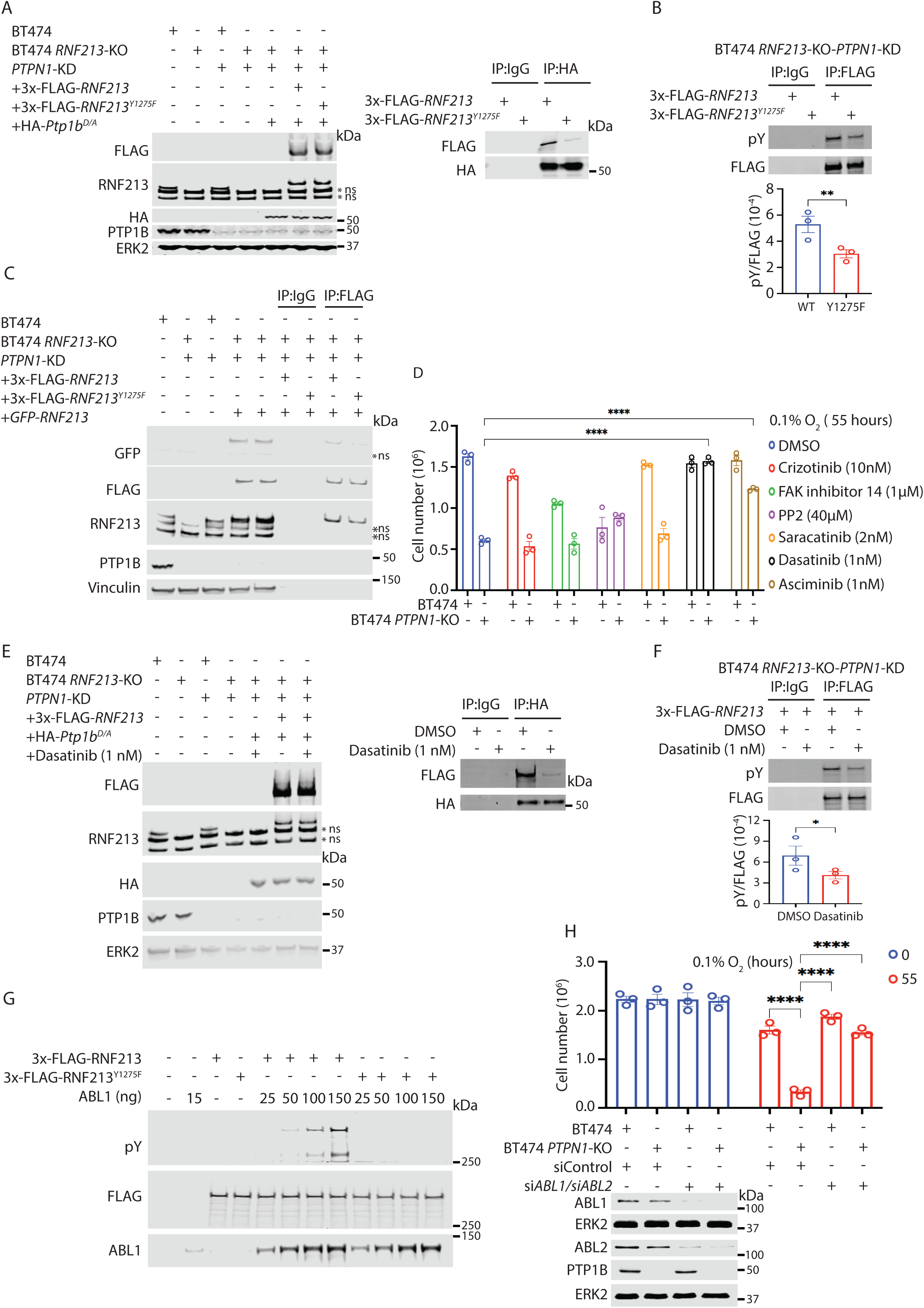
RNF213 phosphorylation at Y1275 is regulated by PTP1B and ABL1/2 and controls oligomerization. (a) RNF213 Y1275 phosphorylation regulates interaction with PTP1B substrate-trapping mutant. BT474 *RNF213*-KO-*PTPN1*-KD cells, with or without stable expression of a doxycycline (Dox)-inducible, HA-tagged substrate trapping mutant (D181A) of mouse PTP1B (HA-*Ptpn1^D/A^*), were transiently transfected with 3x-FLAG-*RNF213* or 3x-FLAG-RNF213^Y1275F^ and exposed to 0.5 μg/ml Dox (+) or left untreated (-). Lysates (left) were subjected to anti-HA immunoprecipitation and immunoblotting for the indicated proteins (right). ERK2 serves as a loading control. (b) Y1275 is phosphorylated in BT474 cells. BT474 *RNF213*-KO-*PTPN1*-KD cells were transiently transfected with 3x-FLAG-*RNF213* or 3x-FLAG-*RNF213^Y^*^1275^*^F^*. Lysates were subjected to anti-FLAG immunoprecipitation and immunoblotting, as indicated, and pY/FLAG signal (each in arbitrary units) was quantified. (c) Y1275 phosphorylation promotes RNF213 oligomerization. BT474 *RNF213*-KO-*PTPN1*-KD cells were transiently transfected with *GFP*-*RNF213* and 3x-FLAG-*RNF213* or 3x-FLAG-*RNF213^Y^*^1275^*^F^*. Lysates were subjected to anti-FLAG immunoprecipitation/immunoblotting. Vinculin serves as a loading control. (d) ABL kinases promote RNF213-mediated hypoxia hypersensitivity. BT474 and BT474 *PTPN1*-KO cells were treated with the indicated kinase inhibitors. Cells were subjected to 0.1 % O_2_ for 55 hours, and viable cells were counted. Error bars represent ± S.E.M.. (e) Dasatinib inhibits RNF213-PTP1B interaction. BT474 *RNF213*-KO-*PTPN1*-KD cells expressing substrate trapping mutant HA-*Ptpn1*^D/A^ were transiently transfected with 3x-FLAG-*RNF213* and then treated with dasatinib or vehicle (DMSO) for 24 hours. Cell lysates were subjected to anti-HA immunoprecipitation/immunoblotting. (f) Dasatinib inhibits tyrosine phosphorylation of RNF213. BT474 *RNF213*-KO-*PTPN1*-KD cells were transiently transfected with 3x-FLAG-*RNF213* followed by treatment with DMSO or dasatinib for 24 hours, and lysates were subjected to anti-FLAG immunoprecipitation/immunoblotting. (g) ABL phosphorylates RNF213 on Y1275. 3x-FLAG-RNF213 and 3x-FLAG-RNF213^Y1275F^ were purified and phosphorylated *in vitro* with the indicated amounts of recombinant ABL1 (see Methods) (h) ABL1/ABL2 promotes RNF213 mediated hypoxia hypersensitivity in *PTPN1*-KO cells. BT474 and BT474 *PTPN1*-KO cells were transfected with *ABL1* and/or *ABL2* siRNAs targeting or control non-targeting siRNA. Cells were subjected to 0.1% O_2_ for 55 hours, and viable cells were counted (top). Error bars represent ± S.E.M. Bottom panel shows immunoblots for the indicated proteins. All experiments (a-g) are representative results from three independent experiments except for the quantifications in b, d, f, h, which are pooled data from the triplicates. Panels b, f were analyzed by ratio paired two-tailed t-test; all other results were evaluated by two-way ANOVA (analysis of variance), \**P*<0.05, \*\*\*\**P*<0.0001, ns=non-specific band.

PTP1B deficiency/inhibition markedly increases overall cellular ubiquitylation, dependent on RNF213^22^. Y1275 is distant from the RING or RZ domains in the monomeric RNF213 structure^23^, arguing against direct regulation. We therefore tested the hypothesis that phosphorylation affects RNF213 oligomerization by co-transfecting expression vectors for 3x-FLAG-*RNF213* or 3x-FLAG-*RNF213^Y^*^1275^*^F^* and *GFP-RNF213* and performing anti-FLAG immunoprecipitations. Notably, recovery of GFP-RNF213 in 3x-FLAG-RNF213^Y1275F^ immunprecipitates was impaired in BT474 (Fig. 1c) and AU565 cells (Extended Data Fig. 1e).

We next sought the kinase(s) that phosphorylate(s) RNF213-Y1275. ABL family kinase inhibitors (dasatinib, asciminib) abrogated the hypoxia hypersensitivity of BT474 *PTPN1*-KO cells and SKBR3 *PTPN1*-KO cells (Fig. 1d; Extended Data Fig. 2a). Dasatanib treatment (compared to treatment with DMSO vehicle alone) also dramatically reduced the recovery of RNF213 in PTP1B trapping mutant immunoprecipates from these cells (Fig. 1e, right; Extended Data Fig. 2b) and reduced overall RNF213 tyrosine phosphorylation to an extent similar to that caused by Y1275F mutation (Fig. 1f; Extended Data Fig. 2c). We therefore asked whether ABL1 phosphorylates RNF213 *in vitro*. We immunopurified FLAG 3x-FLAG-RNF213 and 3x-FLAG-RNF213^Y1275F^ (Extended Data Fig. 2d), incubated them with recombinant, His-tagged ABL1, and found that His-ABL1 phosphorylated WT-RNF213, but not RNF213^Y1275F^, in a ABL1-concentration and time -dependent manner (Fig. 1g; Extended Data Fig. 2e). Furthermore, *ABL1/2* siRNA-treated cells were resistant to hypoxia hypersensitivity induced by PTP1B deficiency (Fig. 1h). Similar results were observed using parental and *PTPN1*-KO MDAMB361 cells (Extended Data Fig. 2f). We conclude that RNF213 Y1275 phosphorylation is regulated by PTP1B and ABL1/2, and phosphorylation of this site promotes RNF213 oligomerization and hypoxia hypersensitivity.

### RNF213 interacts with CYLD and SPATA2

To begin to understand how RNF213 regulates hypoxia sensitivity, we identified RNF213-interacting proteins (Fig. 2a). TurboID-*RNF213* (“N-Turbo”), *RNF213*-TurboID (“C-Turbo”), or TurboID alone (Control Turbo) were recombined into HeLa FlpIn-TRex *RNF213*-KO cells. These cells were labelled with biotin for 10 min., and biotinylated proteins were recovered on streptavidin, digested with trypsin, and analyzed by MS (Fig. 2a). Proteins were ranked as potential binding partners by using the SAINT express algorithm^32^. Immunoblotting (Extended Data Fig. 3a) and silver staining (Extended Data Fig. 3b) confirmed that similar amounts of each sample were submitted for analysis. Relatively few proteins were labelled by N-Turbo (Extended Data Fig. 3c). By contrast, C-Turbo labelled PTP1B (PTPN1) and USP15 (blue-colored arrows), previously reported to interact with RNF213,^22,33^ as well as several new proteins were significantly enriched over the control sample (Fig. 2b, Supplementary Table 2). Among the other top candidates were the major K63-deubiquitinase and NF-κB regulator CYLD and its interacting protein SPATA2 (Fig. 2b, orange-colored arrows). RNF213/CYLD interaction was recently reported by another group, although its biological significance was not determined^34^. Other top C-Turbo interactors included proteins involved in VEGF signaling, which is induced in hypoxia (FLII, MYOF), inflammation (MAP4K4, SPTLC1), lipotoxicity (SPTLC1, ALDH1B1), and ischemia response (CPEB4),^35,36,37,38,39,40,41^ all of which could be potentially relevant for hypoxic cancers. Top N-Turbo interactors included CYP51A1, which plays a crucial role in cholesterol metabolism and atherosclerosis^42^. Pathway analysis suggests involvement of RNF213 in K63-linked deubiquitylation, cytokine (including interferon)-mediated signaling, innate immunity, necroptotic cell death, and cytoskeletal regulation (Extended Data Fig. 3d, 3e).

**Figure 2.**
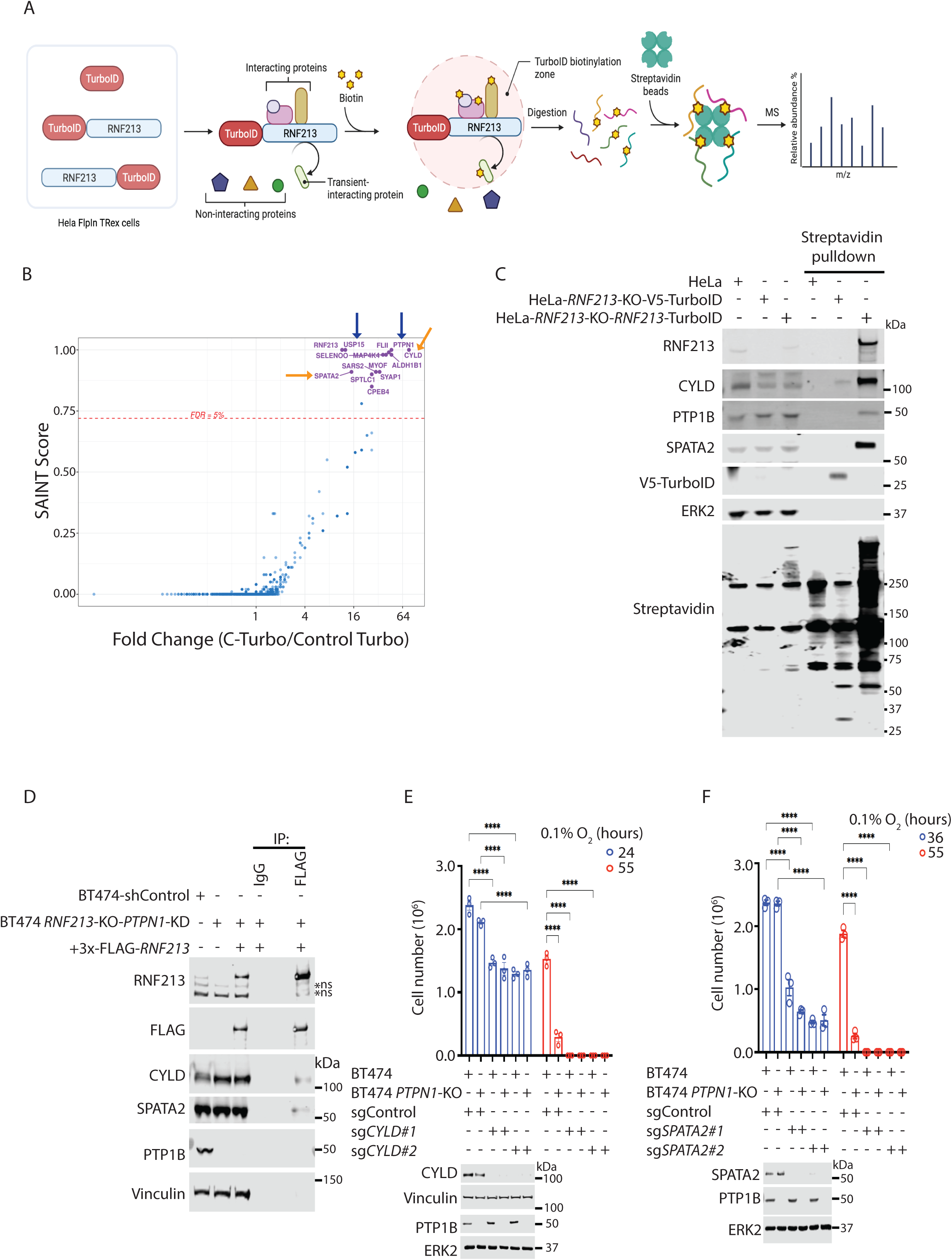
RNF213 interacts with CYLD/SPATA2. (a) Schematic showing proximity-based labelling of RNF213 interactors. HeLa FlpIn-TRex cells expressing TurboID (control), TurboID-RNF213 (N-Turbo) or RNF213-TurboID (C-Turbo) were used for proximity-based biotin labeling. (b) One-sided volcano plot showing the fold-change and SAINT score of proteins identified in C-Turbo versus Control Turbo. Proteins with a SAINT Score >5% FDR are highlighted and considered to be potential interacting partners of RNF213. (c) CYLD and SPATA2 are in proximity with RNF213. Lysates prepared as in Fig. 2a were subjected to streptavidin affinity purification and immunoblotting for the indicated proteins. (d) CYLD and SPATA2 interact with RNF213. BT474 *RNF213*-KO-*PTPN1*-KD cells were transiently transfected with 3x-FLAG-*RNF213*, and lysates were subjected to immunoprecipitation and immunoblotting as indicated. (e, f) CYLD (e) and SPATA2 (f) protect against hypoxia hypersensitivity. BT474 and BT474 *PTPN1*-KO cells, with or without CYLD, were subjected to 0.1% O_2_ for indicated times, and viable cells were counted (top). Immunoblots (bottom) show absence of CYLD/SPATA2 in stable pools generated with the indicated sgRNAs. Error bars represent ± S.E.M. b, e, f are pooled results from three biological replicates; all other data are representative of these replicates, \*\*\*\**P*<0.0001, two-way ANOVA.

Given their role in NF-κB response, inflammation, and innate immunity, we focused on the CYLD/SPATA2 interaction with RNF213. Immunoblotting confirmed the interaction of C-Turbo, but not Turbo alone, with CYLD, SPATA2, and, as expected, PTP1B; neither Turbo protein interacted with ERK2, which served as a control for non-specific biotinylation (Fig. 2c). CYLD and SPATA2 also co-immunopreciptated with 3x-FLAG-RNF213 expressed by transient transfection of BT474-*RNF213*-KO-sh*PTPN1* cells (Fig. 2d). We next used CRISPR/Cas9 technology and two independent sgRNAs to delete each gene in BT474 and BT474 *PTPN1*-KO cells. Remarkably, *CYLD-*KO or *SPATA2-*KO sensitized BT474 cells to hypoxia even more than *PTPN1-*KO alone (Fig. 2e, 2f). Similar results were obtained when analogous SKBR3-derived cell lines were studied (Extended Data Fig. 4a, 4b). Hence, RNF213 interacts with CYLD and SPATA2, and CYLD and SPATA2 (and, most likely, the CYLD/SPATA2 complex) prevent hypoxia hypersensitivity in breast cancer cell lines.

### RNF213 uses its RZ domain to ubiquitylate CYLD and SPATA2

To test whether CYLD and SPATA2 are RNF213 substrates, we co-expressed ALFA-tagged SPATA2 or CYLD and HA-Ubiquitin (HA-UB) in parental or *RNF213*-KO BT474 cells, and treated these cells with claramine (10μM) or vehicle (Fig. 3a). As expected, co-transfection of UB resulted in an increase in HA-reactive species, which were enhanced further in ALFA-CYLD-or ALFA-SPATA2-expressing cells. Comporting with our previous report^22^, claramine treatment led to an increase in overall ubiquitylation (Fig. 3a, left). Recovery of each ALFA-tagged protein, followed by anti-HA immunoblotting, revealed RNF213-dependent ubiquitylation of CYLD and SPATA2, which was enhanced by PTP1B inhibition (Fig. 3a, right). Analogous results were obtained using AU565 cells (Extended Data Fig. 4c).

**Figure 3.**
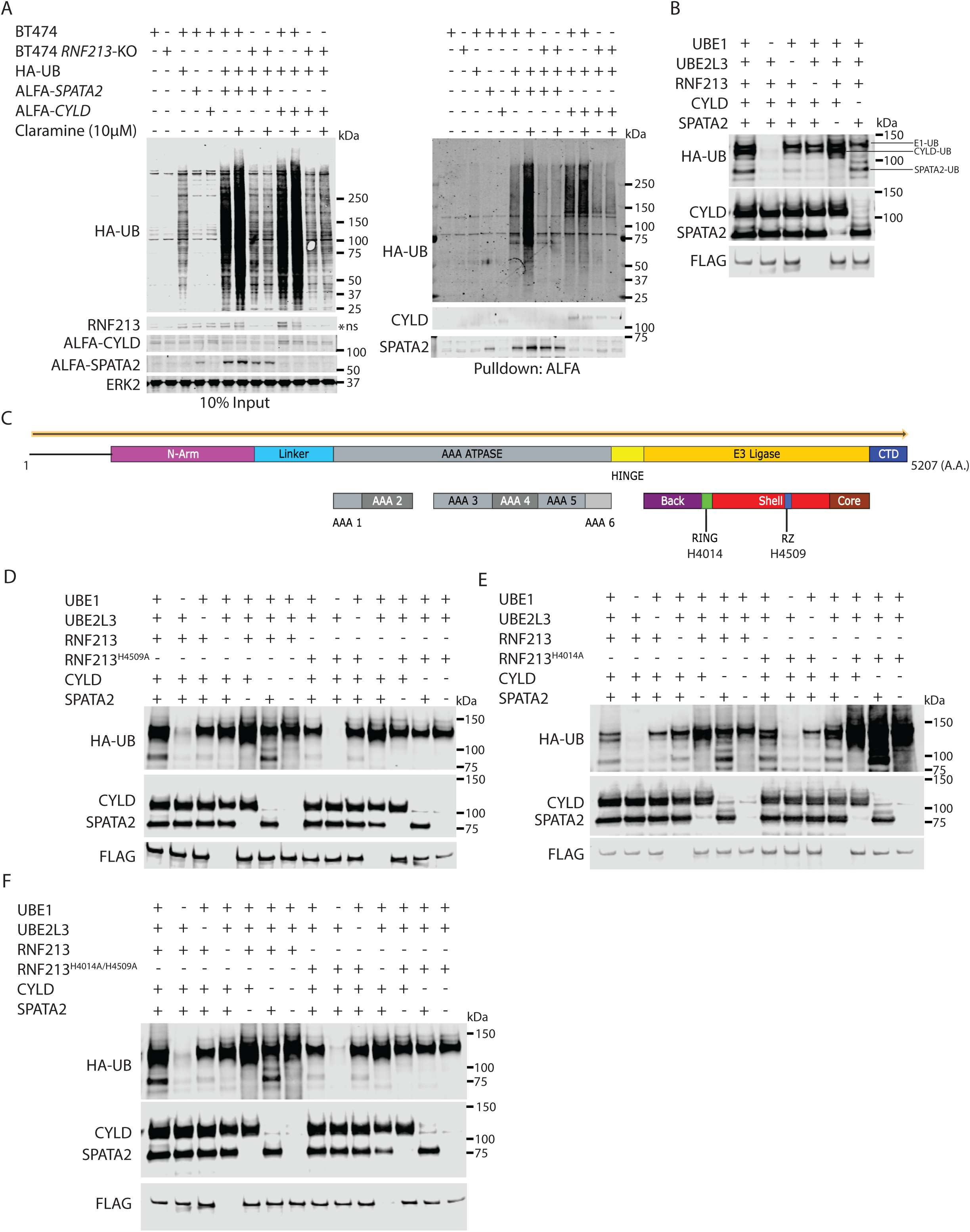
Role of RNF213 RZ and RING domains in CYLD and SPATA2 ubiquitylation. (a) RNF213 promotes SPATA2 and CYLD ubiquitylation. BT474 and BT474 *RNF213*-KO cells were co-transfected with expression contructs for HA-ubiquitin (HA-UB) and ALFA-*SPATA2* or ALFA-*CYLD* and treated with claramine or vehicle (-), as indicated. Lysates (left) were subjected to anti-ALFA immunoprecipitation/immunopblotting (right). (b) *in vitro* ubiquitylation assay shows RNF213/UBE2L3 promotes SPATA2 and CYLD ubiquitylation. Purified wild type was added to *in vitro* ubiquitylation assays with UBE2L3, CYLD, and/or SPATA2 and immunoblotted, as indicated. (c) Domain map of RNF213. Critical residues for RING (H4014) and RZ (H4509) domains are indicated. (d-f) Opposing actions of RNF213 RZ and RING domains in CYLD and SPATA2 ubiquitylation. Purified wild type or mutant 3x-FLAG-RNF213 was added to *in vitro* ubiquitylation assays with UBE2L3, CYLD, and/or SPATA2 and immunoblotted, as indicated. Each blot is representative of two biological replicates, ns=non-specific band.

We purified 3xFLAG-tagged RNF213 and various RNF213 mutants from HeLa FlpIn-TRex *RNF213*-KO cells (Extended Data Fig. 4d top, bottom), and assessed the ability of each protein to ubiquitylate CYLD and SPATA2 *in vitro* using UBE1 as the E1 enzyme and UBE2L3 as the E2. Consistent with our transfection studies, SPATA2 and CYLD, when tested alone or in equimolar concentrations, were ubiquitinated by 3xFLAG-RNF213 (Fig. 3b). As noted above, RNF213 has two potential ubiquitin ligase domains (RING, RZ; catalytic histidines indicated in the cartoon in Fig. 3c). There was no significant ubiquitylation of CYLD or SPATA2 by an RZ mutant (“RZ-dead”) form of RNF213 (RNF213^H4509A^, Fig. 3d). Surprisingly, however, CYLD or SPATA2 ubiquitylation was *increased* when RING mutant RNF213 (RNF213^H4014A^, “RING-dead”) was assayed (Fig. 3e). Based on these results, we hypothesized that the RING negatively regulates the RZ domain. We therefore compared the ubiquitylation activity of WT RNF213 and RNF213^H4014A/H4509A^ (RING*/*RZ-dead). Consistent with our hypothesis RING/RZ-dead RNF213 was unable to ubiquitylate CYLD/SPATA2 (Fig. 3f). Therefore, RNF213 is an E3-ubiquitin ligase for CYLD and SPATA2, their ubiquitylation is catalyzed by the RZ domain, and the RING can restrain RZ activity. Furthermore, RING-dead (or RING-impaired) mutations are RNF213 RZ gain-of-function mutants; notably, highly penetrant MMD SNPs map to the RING domain.

These findings were surprising, because we reported earlier that RING-dead RNF213 or highly penetrant MMD SNPs *decrease* global ubiquitylation in HeLa or 293T cells and appear to act as dominant negative alleles. By contrast, our new data suggest that RING-dead RNF123 is an RZ gain-of-function mutation, which might be expected to *increase* global ubiquitylation. In an attempt to resolve this apparent discrepancy, we co-expressed 3xFLAG-RNF213 or RNF213 mutants with HA-Ubiquitin (HA-UB) in HeLa Flp-In-TRex *RNF213*-KO cells (Extended Data Fig. 4e) and subjected these cells to normoxia or hypoxia (0.1% O_2_ for 24 hours). Consistent with our previous report, we observed a sharp decrease in *global* ubiquitylation with the RING-dead mutant; however, global ubiquitylation also was decreased in cells expressing RZ-dead or RING-*and* RZ-dead RNF213. Similar results were observed in experiments with BT474 *RNF213-*KO cells reconstituted with *RNF213* or various *RNF213* mutants (Extended Data Fig. 4f). These data suggest that RING and RZ domain each have unique substrates, and that negative regulation of the RZ domain by the RING is probably substrate-specific. Alternatively, the RZ domain could regulate targets that, in concert, perturb global ubiquitylation.

### The RNF213 RZ and ATPase domains promote hypoxia hypersensitivity via degradation of SPATA2 and CYLD

To determine which RNF213 domains are required for hypoxia hypersensitivity, we reconstituted BT474 *RNF213*-KO (and AU565-*RNF213*-KO) cells with WT *RNF213*, *RNF213^Y^*^1275^*^F^*, *RNF213^K^*^2775^*^A^* (second AAA-ATPase domain mutant), *RNF213^H^*^4014^*^A^* (RING-dead), *RNF213^H^*^4509^*^A^* (RZ-dead), *RNF213^H^*^4014^*^A/H^*^4509^*^A^* (RING/RZ-dead) or *RNF213^K^*^2775^*^A/H^*^4014^*^A^* (ATPase/RZ-dead), as indicated in cartoon in Fig. 4a. We also used CRISPR/Cas9 to generate *PTPN1*-KO pools of each cell line. The resultant cell lines/pools were subjected to hypoxia, and lysates were analyzed by immunoblotting. As expected, WT RNF213, but not RNF213^Y1275F^, promoted CYLD and, to a lesser extent, SPATA2 degradation in BT474 (Fig. 4b) and AU565 cells (Extended Data Fig. 5a). Also as expected, degradation was increased by PTP1B deficiency. RZ-dead or RING/RZ-dead RNF213 were unable to promote CYLD or SPATA2 degradation, whereas compared with WT RNF213, the RING-dead mutant enhanced degradation (Fig. 4c; Extended Data Fig. 5b), especially in the *PTPN1*-KO background. Previous work showed that RNF213 ATPase activity is required for RZ domain-mediated ubiquitin ligase activity^26^. Given our finding that RING-dead mutations show RZ domain gain-of function, we hypothesized that an active ATPase domain should also be required for substrate degradation by RING-dead mutants. Indeed, a compound ATPase/RING domain mutant (RNF213^K2775A/H4014A^) was unable to degrade CYLD or SPATA2 in the presence or absence of PTP1B (Fig. 4d; Extended Data Fig. 5c). Single nucleotide polymorphisms (SNPs) in *RNF213* are associated with Moyamoya disease (MMD), a rare syndrome characterized by precocious carotid artery stenosis and stroke, often in teenagers or young adults^43–45^. We also tested the effects of select MMD SNPs that affect the RING domain of RNF213,^43,46,47^ by restoring doxycycline-inducible expression of *RNF213^C^*^3997^*^Y^* (highly penetrant) and *RNF213^R^*^4810^*^K^* (most common, low penetrance) to *RNF213*-KO BT474 and AU565 cells. As predicted, the highly penetrant, RING-mutant SNP had similar effects on SPATA2 and CYLD to RING-dead RNF213 (Extended Data Fig. 6a, 6b).

**Figure 4.**
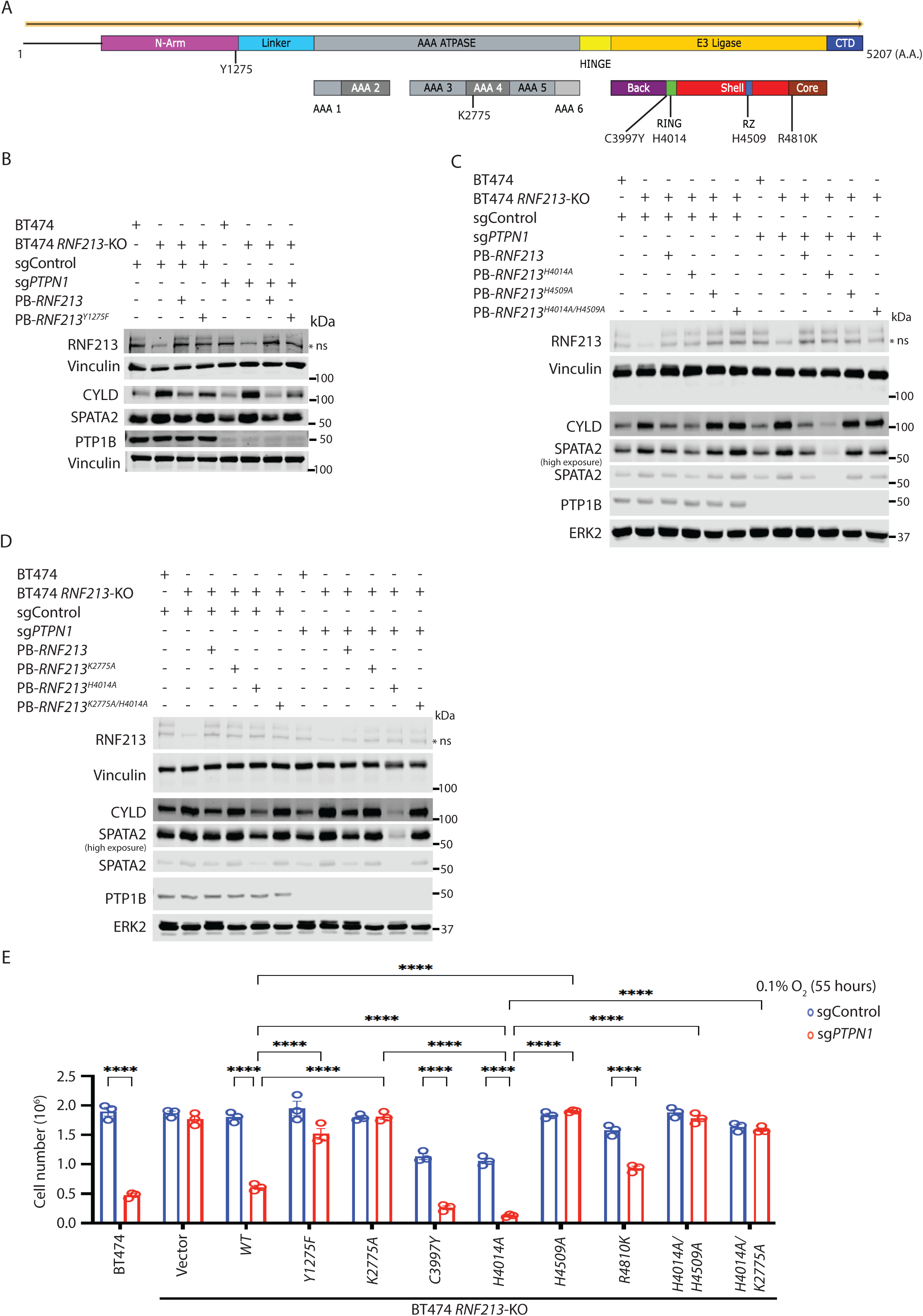
RNF213 RING mutants/highly penetrant MMD SNPs activate RZ domain and require ATPase activity. (a) Schematic showing key domains in RNF213. Critical residues that regulate ATPase (Y1275, K2775), RING, and RZ domains, as well as the positions of select MMD SNPs, are indicated. (b-d) BT474 *RNF213*-KO cell reconstituted with RNF213^WT^, the indicated RNF213 mutants, parental BT474 cells, and BT474 *RNF213*-KO cells with or without PTP1B, were subjected to 0.1% O_2_ for 40 hours. Immunoblotting was performed for the indicated proteins. (e) RZ and ATPase domains of RNF213 are required for hypoxia hypersensitivity of BT474 *PTPN1*-KO cells. Cells prepared as in b-d and Extended Data Fig. 6a were subjected to 0.1% O_2_, and viable cells were counted after 55 hours. Error bars represent ± S.E.M. (n = 3 biological replicates, each panel) and each blot is representative of two biological replicates. \*\*\*\**P*<0.0001, two-way ANOVA, ns=non-specific band.

Finally, we tested the effect of each mutant on hypoxia sensitivity in the presence or absence of *PTPN1*-KO. Consistent with the above biochemical results, cells reconstituted with tyrosine phosphorylation site-, ATPase-, RZ-, compound ATPase/RZ-, or compound RING/RZ-, mutants failed to show hypoxia hypersensitivity in response to PTP1B deficiency (Fig. 4e; Extended Data Fig. 6c). By contrast, RING-dead RNF213 showed increased hypoxic cell death even in PTP1B-replete cells and almost no survival upon *PTPN1* deletion. In concert, though, these data indicate that tyrosine phosphorylation of RNF213 promotes ATPase-mediated oligomerization and activation of the RZ domain ubiquitin ligase activity, which in turn leads to CYLD/SPATA2 ubiquitylation, degradation, and cell death. The RING domain inhibits the RZ domain, and RING-dead RNF213 promote increased CYLD/SPATA2 degradation and hypoxia hypersensitivity.

### The LUBAC complex also is required for CYLD/SPATA2 degradation by RNF213

Recently, it was reported that the RZ domain ubiquitylates *Salmonella* lipopolysaccharide, most likely on its lipid component.^26^ The LUBAC complex then catalyzes further ubiquitylation, promoting bacterial clearance via autophagy. Knockdown of the LUBAC complex component *RBCK1* prevented WT-RNF213-catalyzed degradation of CYLD and SPATA2 even in PTP1B-depleted BT474 or AU565 cells (Fig. 5a; Extended Data Fig. 7a). Remarkably, however, the LUBAC complex was dispensable for CYLD/SPATA2 degradation by RING-dead RNF213. Consistent with these findings, *RBCK1* knockdown was unable to prevent hypoxia-hypersensitivity in *RNF213*-KO cells reconstituted with the RING-dead mutant (Fig. 5b, Extended Data Fig. 7b).

**Figure 5.**
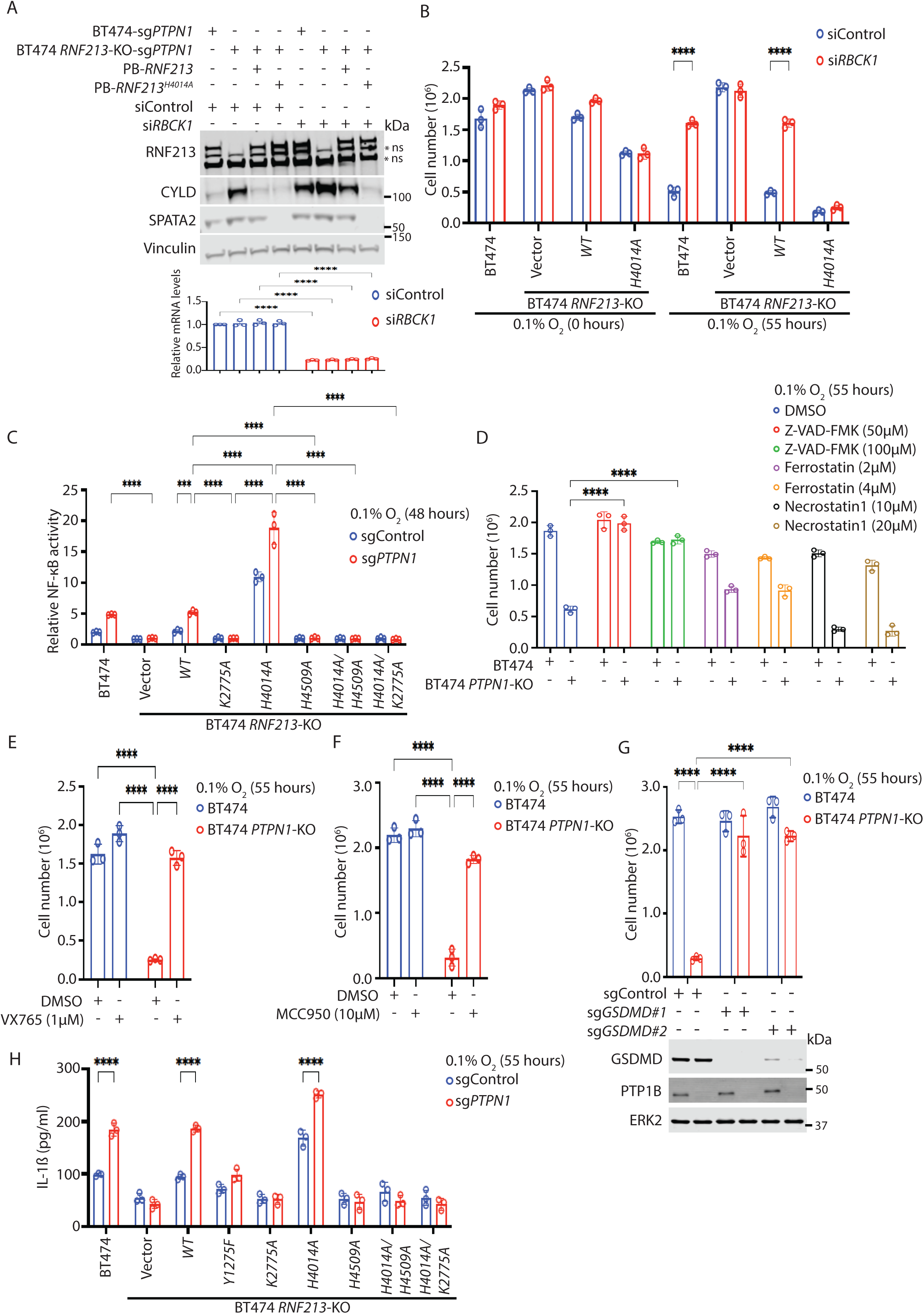
*PTPN1*-KO cells die via RNF213-and LUBAC-dependent pyroptosis. (a) Wild-type, but not RING-dead RNF213 requires LUBAC for CYLD/SPATA2 degradation. BT474 *RNF213*-KO cells reconstituted with wild-type *RNF213* or *RNF213^H^*^4014^*^A^* parental BT474 cells and BT474 *RNF213*-KO with or without PTP1B were transfected with *RBCK1* or control siRNA. Cells were subjected to 0.1% O_2_ for 40 hours, lysed, and, immunoblotted as indicated (top). *RBCK1* knockdown was confirmed by RT-qPCR (bottom). (b) RING-dead RNF213 bypasses LUBAC requirement. The indicated cells were subjected to 0.1% O_2_ for 55 hours, and viable cell numbers were counted. (c) RNF213 promotes NF-κB activity. Cells described in Fig. 4e were transfected with an NF-κB reporter construct (see Methods) and subjected to 0.1% O_2_ for 48 hours. Firefly and Renilla luciferase signals were measured, reporter activity was quantified as the ratio of Firefly/Renilla signal, and all other signals were normalized to that from parental BT474 cells. (d) Hypoxic cell death in *PTPN1*-KO cells is caspase-dependent. BT474 and BT474 *PTPN1*-KO cells were treated with the indicated inhibitors, placed in 0.1% O_2_ for 55 hours, and cell viability was assessed. (e, f) Hypoxic cell death in *PTPN1*-KO cells requires inflammatory caspases and the inflammasome. As in (d), except with/without VX765 (e) or MCC950 (f). (g) GSDMD promotes hypoxic cell death of *PTPN1*-KO cells. Pools of BT474 and BT474 *PTPN1*-KO cells, with or without *GSDMD* deletion, were subjected to 0.1% O_2_, and cell viability was measured at 55 hours (top). Immunoblot indicates efficiency of *GSDMD* knockout (bottom). (h) Hypoxia-hypersensitive cells release IL-1β. Cells prepeared as in Fig. 4e were subjected to 0.1% O_2_ for 55 hours, and IL-1β in supernatants was quantified by immunoassay. In all panels, error bars represent ± S.E.M. (n = 3 biological replicates) and each blot is representative of two biological replicates.\*\*\*\**P*<0.0001, two-way ANOVA, ns=non-specific band.

### RNF213, via its RZ domain, promotes NF-κB activation and primes cells for pyroptosis

CYLD and SPATA2 are important negative regulators of NF-κB, so CYLD/SPATA2 degradation would be expected to increase NF-κB activity. We used a dual luciferase reporter assay to monitor NF-κB activity in *RNF213*-KO BT474 and AU565 cells reconstituted with WT *RNF213* and various mutants with or without concomitant (pooled) *PTPN1* deletion. *RNF213*-KO cells (Vector) failed to increase reporter activity when PTP1B was depleted, while reconstituting these cells with WT-*RNF213* increased activity in *PTPN1*-KO cells to levels similar to that of parental BT474 cells with *PTPN1* deletion. RING-dead RNF213-reconstituted cells had markedly increased NF-κB activity even when *PTPN1* was intact, and activity increased further upon *PTPN1* deletion. Consistent with their effects on CYLD/SPATA2, RZ mutant, RING/RZ mutant, or ATPase/RZ mutant forms of RNF213 failed to induce NF-κB under any condition and might even have a dominant negative effect (Fig. 5c; Extended Data Fig. 7c).

CYLD promotes apoptosis and necroptosis,^48,49^ and NF-κB typically promotes cancer cell survival^50^. But PTP1B depletion or inhibition increases cell death in hypoxia, while promoting CYLD degradation and enhanced NF-κB activity. To resolve this apparent paradox, we investigated how PTP1B-deficient, hypoxic cells die, initially by testing the effects of known inhibitors of various cell death pathways. Neither ferroptosis nor necroptosis inhibitors blocked hypoxia hypersensitivity in *PTPN1*-KO cells; however, the general caspase inhibitor Z-VAD-FMK prevented hypoxia-induced cell death (Fig. 5d). Similar results were obtained using SKBR3 or MDAMB361 cells (Extended Data Fig. 7d, 7e).

We previously observed no increase in apoptosis (a caspase-dependent process) in PTP1B-deficient breast cancer cells; instead, death appeared to be necrotic^22^. Z-VAD-FMK also inhibits “inflammatory caspases” (Caspases 1,4, and 5 in humans, Caspases 1 and 11 in mouse). Remarkably, treatment with VX765 (1 μM), a specific inhibitor of Caspase 1 and Caspase 4, completely blocked hypoxic cell death in *PTPN1*-KO BT474 cells (Fig. 5e). Similar results were observed using MDAMB361 cells (Extended Data Fig. 7f).

NF-κB induces the expression of the inflammasome component *NLRP3*, as well as *IL1B*. The inflammasome, in turn, facilitates pro-IL-1β cleavage and release of the active cytokine, as well as the cleavage of Gasdermins^51^. Cleaved Gasdermins oligomerize and form pores in cell membranes, leading to pyroptotic cell death. Notably, hypoxia hypersensitivity of *PTPN1*-KO BT474 or MDAMB361 cells was also blocked by treatment with the NLRP3 inhibitor MCC950 (Fig. 5f; Extended Data Fig. 7g) or by knockout of *GSDMD* with either of two sgRNAs (Fig. 5g; Extended Data Fig. 7h). *PTPN1*-KO also increased IL-1β production in RNF213-WT-reconsituted BT474 or AU565 cells (Fig. 5h; Extended Data Fig. 7i). IL-1β levels increased further in cells expressing RING-dead RNF213, but there was no detectable increase in cells expressing RNF213 that cannot be tyrosine phosphorylated (Y1275F) or in RZ-dead, ATPase-dead, RING-dead and RZ-dead or RING-dead and ATPase-dead mutants.

### Hypoxia-driven ER stress sends the second signal for pyroptotic cell death

NF-κB can prime cells for pyroptosis, but a “second signal” is required to trigger inflammasome assembly/inflammatory caspase activation, Gasdermin cleavage, and cell death. Several stimuli can provide the second signal, including ER stress, which is evoked by O_2_ levels ≤1%^52^. To address the relevance of O_2_ levels and its connection with ER stress, we assessed the viability of *PTPN1*-KO cells with and without CYLD or SPATA2 (Extended Data Fig. 8a). *PTPN1*-KO or depletion of CYLD and SPATA2 sensitized BT474 cells to death 0.1% - 0.2% O_2_ (Extended Data Fig. 8a). At 0.5% O_2_, combined *PTPN1/CYLD*-KO, *PTPN1/SPATA2*-KO, or *CYLD-*KO alone, showed increased hypoxia sensitivity. There were no significant effects of any of these mutants on viability in 1% O_2_. Similar results were obtained when analogous MDAMB361-derived cell lines were studied (Extended Data Fig. 8b). Remarkably, *PTPN1*-KO cells treated with the IRE1α inhibitor 4μ8c were resistant to hypoxic cell death compared with DMSO control-treated cells (Fig. 6a). Similar results were obtained using MDAMB361 cells (Extended Data Fig. 8c) or by treating *PTPN1*-KO BT474 or MDAMB361 cells with the PERK inhibitor GSK2606414 (Fig. 6b; Extended Data Fig. 8d). These data show that ER stress signaling is required for hypoxia sensitivity in PTP1B-deficient cells.

**Figure 6.**
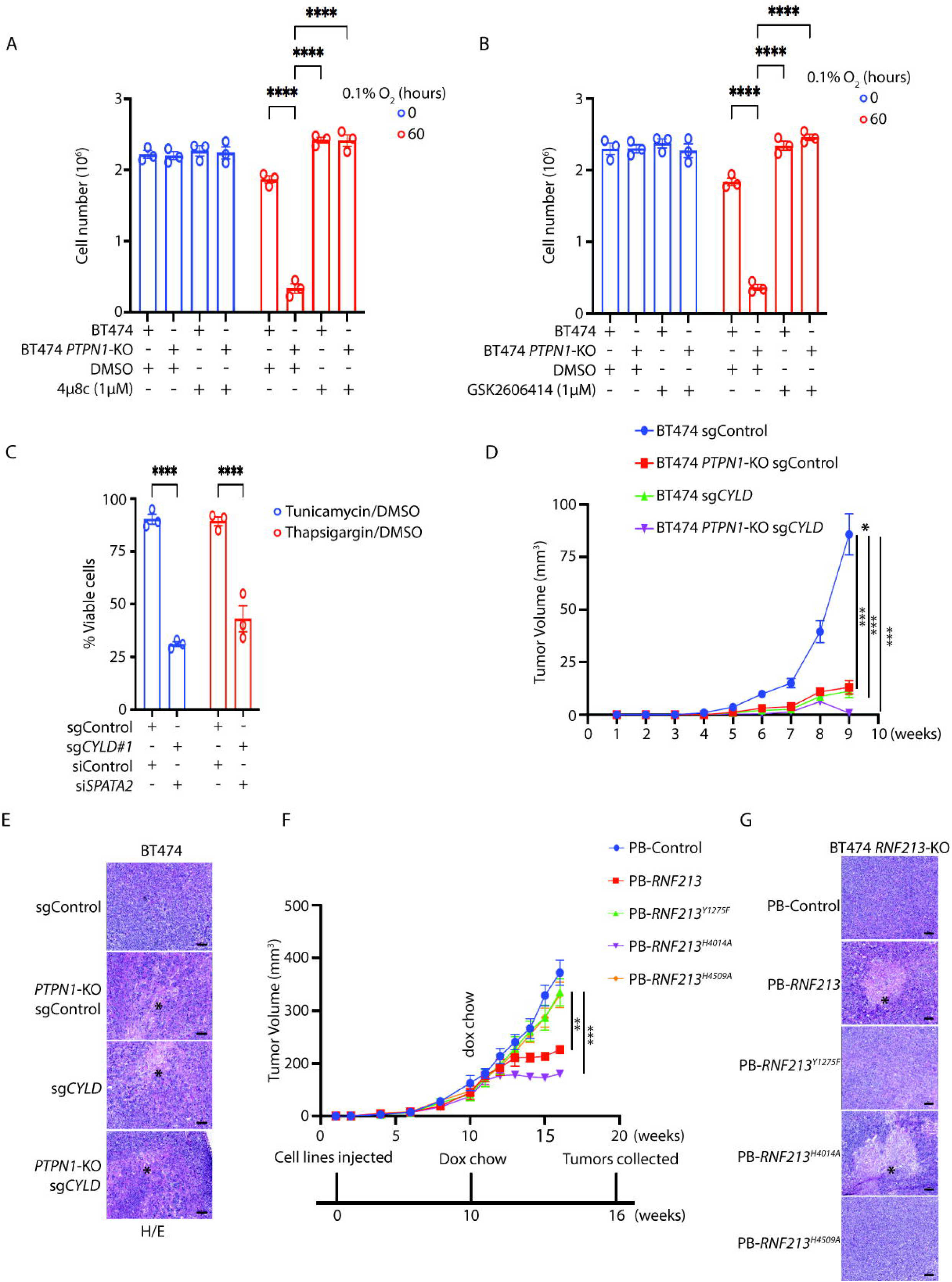
Hypoxia-evoked ER stress activates the inflammasome and CYLD controls HER2+ breast cancer tumorigenesis. (a, b) ER stress promotes hypoxia hypersensitivity. BT474 and BT474 *PTPN1*-KO cells treated with the IRE1α inhibitor 4μ8C (a) or the PERK inhibitor GSK2606414 (b), were subjected to 0.1% O_2_ for 55 hours, and viability was assessed. (c) ER stress causes pyroptosis in CYLD/SPATA2-deficient cells in normoxia. BT474 cells, with or without CYLD/SPATA2, were treated with tunicamycin (10 μM) or thapsigargin (5 μM) for 40 hours, and cell viability was measured at 40 hours. (d, e) CYLD is critical for HER2+ breast tumorigenesis. BT474 and BT474 *PTPN1*-KO cells, with or without *CYLD*-KO, were injected subcutaneously into athymic *nu/nu* mice (6/group). Tumor size was measured weekly for 9 weeks (d). Residual tumors were excised, and fixed and stained with H&E (e). * in (e) indicates representative necrotic regions. (f, g) RNF212 RZ domain and tyrosine phosphorylation site are critical for HER2+ breast tumorigenesis. BT474 *RNF213*-KO sg*PTPN1* cells were reconstituted with vectors expressing WT *RNF213*, *RNF213^Y^*^1275^*^F^* (phosphosite-mutant), *RNF213^H^*^4014^*^A^* (RING-dead), or *RNF213^H^*^4509^*^A^* (RZ-dead) and implanated into athymic *nu/nu* mice (6/group). When tumors were of significant size, mice were switched to doxycycline chow to induce expression of WT or mutant RNF213. Tumor size was measured weekly (f). Residual tumors were excised, and fixed and stained with H&E (g). * in (g) indicates representative necrotic regions. Error bars represent ± S.E.M., \*\*\**P*<0.001, two-way ANOVA.

To ask whether ER stress is sufficient to send the second signal, we treated parental BT474 cells and BT474 cells rendered SPATA2-and CYLD-deficient (by *CYLD*-KO and *SPATA2* siRNA) to mimic the effects of PTP1B deficiency in hypoxia with tunicamycin or thapsigargin in normoxia. Both agents induced cell death selectively in SPATA2-and CYLD-deficient cells (Fig. 6c). Analogous results were obtained using cognate MDAMB361 cells (Extended Data Fig. 8e).

### CYLD levels are critical for HER2+ tumor growth

To explore the potential role of CYLD in human HER2+ BC tumorigenesis, we implanted parental BT474, BT474 *PTPN1*-KO, BT474-*CYLD*-KO, and double knockout cells into the flanks of athymic *nu/nu* mice and monitored tumor development. The absence of either PTP1B or CYLD (or both) markedly inhibited tumor growth (Fig. 6d). Similar results were obtained with analogous MDAMB361-derived cell lines (Extended Data Fig. 9a). Histological staining (H&E) of tumors formed with BT474 cells showed increased necrosis in *PTPN1*-KO, *CYLD*-KO or *PTPN1*/*CYLD*-KO compared with those from parental cells, even though the former were much smaller (Fig. 6e). Similar results were obtained with analogous MDAMB361-derived cell lines (Extended Data Fig. 9b). To determine which RNF213 domains were required for tumor death, we implanted BT474 *RNF213*-KO sg*PTPN1* cells reconstituted with doxycycline-inducible WT *RNF213*, *RNF213^Y^*^1275^*^F^*(phosphosite-mutant), *RNF213^H^*^4014^*^A^* (RING-dead), or *RNF213^H^*^4509^*^A^* (RZ-dead). When tumors were of significant size, mice were switche to doxycycline chow to induce the expression of WT or mutant RNF213. As expected, expression of WT, as well as RING-dead, RNF213 impaired tumor growth; by contrast, tumors expressing phosphosite or RZ-dead mutant-reconstituted cells behaved like RNF213-KO tumors (Fig. 6f). Despite their decreased size, WT *RNF213*-and *RNF213^H^*^4014^*^A^*-reconstituted tumors also showed marked necrosis (Fig. 6g).

## Discussion

We have uncovered a unique mechanism of inflammatory cell death in breast cancer cells exposed to hypoxia (Extended Data Fig. 9c). We find that RNF213, the product of the major susceptibility gene for MMD, is phosphorylated at Y1275. Y1275 phosphorylation promotes RNF213 oligomerization, and is reciprocally regulated by ABL1,2 and PTP1B. Oligomerization activates the RNF213 RZ domain to catalyze CYLD/SPATA2 ubiquitylation and degradation. The LUBAC complex also is required for RNF213-initiated degradation of CYLD/SPATA2, but this requirement is abrogated in MMD SNPs or by engineered mutants that inactivate the RNF213 RING. Decreased CYLD/SPATA2 levels lead to increased NF-κB activation in hypoxia, which promotes *NLRP3* transcription. A second signal, most likely hypoxia-induced ER stress, promotes inflammasome assembly and activation, leading to pro-IL-1β processing/secretion and GSDMD-dependent pyroptotic cell death. Previously, it has been reported that PD-L1 expression and its subsequent nuclear localization leads to GSDMC expression in hypoxia that triggers pyroptosis^53^. Our results are independent of PD-L1 as it has been reported that BT474 and other cell lines like MCF7 does not have detectable levels of PD-L1^54–56^ although they are sensitive to hypoxia after PTP1B inhibition or deficiency (Extended Data Fig 1a). Our results, provide new insights into tumor biology and the hypoxia response, particularly the link between hypoxia and inflammation and how these can be exploited for therapy.

NF-κB signaling is activated in hypoxia through IKKβ, but the detailed mechanism has remained unclear. Scholz *et al.*^57^ reported that components of the TRAF6 complex (OTUB, UVEV1a) associate with PHD1 and that multiple NF-κB pathway components are ubiquitylated. Our results indicate that hypoxia also promotes NF-κB activation via degradation of a major deubiquitylase complex. HPV-infected cells activate NF-κB in hypoxia via CYLD degradation.^58^ Intriguingly, degradation requires E6, but not E6AP, the ubiquitin ligase that typically mediates E6 action. It will be interesting to test whether RNF213 is involved in this process.

Most previous studies of RNF213 have focused on its E3-ubiquitin ligase domains, although it is known that its E3 ligase activity requires ATPase function and, most likely, hexamerization^26^. Oligomerized RNF213 serves as a sensor for ISG15^59^, but to our knowledge, reciprocal control of Y1275 phosphorylation by ABL1/2 and PTP1B is the first example of a signaling pathway regulating RNF213 oligomerization/E3 activation. We also find that the RZ domain, previously implicated in lipid ubiquitylation, has protein targets (CYLD/SPATA2). How a single E3 can have such dual activities awaits futher biochemical/structural investigation.

Hypoxic regions in tumors typically are therapy-resistant and constitute a cellular reservoir for tumor recurrence^3,7^. PTP1B inhibition or degradation (e.g., using PROTAC technology)^60^ can kill these hypoxic cells and could be useful in combination with other anti-neoplastics. Importantly, we find that PTP1B inhibition/deficiency induces pyroptosis, an inflammatory form of cell death, which could help “turn cold tumors hot” and promote the anti-tumor immune response^51^. However, inflammation also can suppress tumor immunity^51^; notably, PTP1B inhibition/deficiency increases IL-1β production, which promotes immunosuppression by myeloid cells^61^. PTP1B also has direct effects in myeloid cells^62^ and lymphocytes^63,64^; whether the PTP1B/RNF213/CYLD/SPATA2 pathway is active in these cells remains unclear. PTP1B/TCPTP inhibitors are in clinical trials as anti-neoplastic agents^30^ (NCT04777994), mainly because of its predicted immunostimulatory effects^63,65^. Our results suggest that PTP1B inhibition might also have important, salutary direct effects on tumors cell. Elucidating the cell-autonomous and non-autonomous effects of such agents, including if/how TCPTP inhibition affects RNF213, should aid in their rational deployment. It also will be important to test whether PTP1B controls hypoxia sensitivity in other tumor types, such as pancreas cancer, which is notoriously hypoxic^66^.

## Acknowledgements

This work was funded by RO1 CA049152, RO1 CA264933, and P30 CA016087 to B.G.N. We thank the Perlmutter Cancer Center Proteomics Research Laboratory (PRL) and Experimental Pathology Laboratory (ExPath), supported in part by P30CA016087, for technical support and help with data analysis and interpretation. Mass spectrometric experiments were also supported by NIH shared instrumentation grant 1S10OD010582-01A1. We thank Drs. Leslie Mitchell and Jef D. Boeke from the Institute of System Genetics, NYU Grossman School of Medicine for help with purification of full-length RNF213, supported by CEGS grant 1RM1HG009491.

## Author Contributions

A.B. conceptualized the project, designed and performed all experiments except MS analysis, curated and analyzed the data, and wrote initial drafts of the manuscript. M.A. performed LC MS/MS experiments to identify RNF213-interacting proteins/substrates. B.U. supervised all proteomics experiments. B.G.N. conceptualized the project, provided supervision, funding, and project administration, and wrote/edited the final manuscript.

## Methods

### Cell culture and cell line generation

Sources of the breast cancer cell lines used here were described;^67^ all are maintained as frozen stocks by the Neel Laboratory. Except where indicated, cells were cultured in DMEM +10% FBS with 100 U/ml penicillin and 100 mg/ml streptomycin at 37°C in 5% CO_2_ and tested monthly for mycoplasma by PCR.

Isogenic HeLa Flp-In T-Rex *RNF213*-KO cells that inducibly express wild-type *RNF213* or *RNF213* variants in response to doxycycline (DOX) were generated by co-transfecting pcDNA5/FRT/TO-based expression constructs with 3x-Flag tagged *RNF213* or *RNF213* variant with pOG44, which directs constitutive expression of Flp recombinase (FLP). Cells with successful integration were selected by adding hygromycin B (200 μg/ml) 48 hours post-transfection. Single cell clones of *RNF213*-deleted HeLa, BT474, and AU565 cells, or of *PTPN1-* deleted BT474, SKBR3, and MDAMB361 cells, were generated by using CRISPR/Cas9 technology.^68^ Briefly, sgRNAs targeting each gene were cloned into the BbsI site of pSpCas9 (BB)–2A–Puro (PX459; Addgene), and cells were transfected with each cloned vector using Lipofectamine™ 3000 Transfection Reagent (ThermoFisher Scientific). After 48 hours, cells were diluted into 96-well plates (1 cell/well) containing 48-hour conditioned media from the same line in addition to standard media. Knockout clones were confirmed by immunoblotting. The sgRNA sequences are shown in Supplementary Table 3.

Piggbac (PB)-based expression was used to re-express WT *RNF213* or RNF213 variants under Dox control in *RNF213*-KO BT474 and *RNF213*-KO AU565 cells. Briefly, PB-TA-ERN (Addgene)-based expression constructs containing RNF213 and RNF213 variants were co-transfected with transposase expression vector, and integrants were selected by adding G418 (800μg/ml). RNF213 expression was induced by adding 0.5 μg/ml Dox.

*PTPN1* depletion was accomplished by stable shRNA knockdown using pSUPER-Puro-based retroviral vectors. *RBCK1*, *ABL1*, and *ABL2* ON-TARGETplus siRNA (Horizon Discovery) were introduced using Lipofectamine™ RNAiMAX reagent (ThermoFisher Scientific). Pooled knockout lines were generated by infection with lentiviruses expressing CAS9 and appropriate sgRNAs. Lentiviruses and retroviruses were packaged as described.^69^

### Immunoblotting, immunoprecipitation, and SDS-PAGE

For co-immunoprecipitations, cells were lysed in NP40 buffer (50mM Tris⋅HCl [pH 8], 150mM NaCl, 2mM EDTA, 10mM Na4P2O7, 100mM NaF, 2mM Na3VO4, 1% [vol/vol] NP-40, 40 μg/ml phenylmethyl sulfonyl fluoride, 2 μg/ml antipain, 2μg/ml pepstatin A, 20μg/ml leupeptin, and 20μg/ml aprotinin). For streptavidin affinity purifications and standard immunoprecipitations, modified RIPA buffer (50mM Tris⋅HCl [pH 8], 150mM NaCl, 2mMEDTA, 10mM Na4P2O7, 100mM NaF, 2mM Na3VO4, 1% [vol/vol] NP-40, 0.1% (wt/vol) SDS, 40μg/ml phenylmethyl sulfonyl fluoride, 2μg/ml antipain, 2μg/ml pepstatin A, 20μg/ml leupeptin, and 20μg/ml aprotinin) was used for cell lysis. For detection of ubiquitylated species *in vivo*, 10mM N-ethylmaleimide (NEM) and 10mM iodoacetamide (IAM) were added to the lysis buffer.

For immunoblotting of whole cell lysates, equal amounts of protein per sample were subjected to SDS–PAGE and transferred to polyvinylidene fluoride (PVDF) membranes (Millipore). For immunoprecipitation of FLAG-tagged proteins, lysates were incubated with anti-FLAG-magnetic beads for 4 hours on a rotator-mixer at 4°C. For immunoprecipitation of HA-tagged proteins, lysates were incubated with anti-HA-agarose beads for 4 hours on a rotator-mixer at 4°C. For immunoprecipitation of ALFA-tagged proteins, lysates were incubated with ALFA-selector ST agarose beads overnight on a rotor-mixer at 4°C. For recovery of biotinylated proteins, high-capacity streptavidin agarose resin (ThermoScientific) was added to lysates, which were rotated for 4 hours at 4°C. Beads were washed five times in lysis buffer, incubated in sample buffer at 95C for 5 min. and resolved by SDS-PAGE and immunoblotting. For in vitro ubiquitylation and phosphorylation using purified RNF213 and RNF213 variants, proteins were eluted from beads by incubation with an excess of 3X-FLAG peptide. Lysates and immunoprecipitates were resolved by modified SDS–PAGE on 3–8% Tris-acetate gels (Novex) or 5% Tris-glycine gels. Gels were transferred in 1× transfer buffer, 10% methanol for 1 hour at 25 V using an XCell II Blot Module (Novex) and analyzed by acquisition software (Image Studio Lite), on an Odyssey Infrared imaging system (Li-Cor Biosciences). All antibodies used are listed in Supplementary Table 3.

### Viability assays

Cells were seeded in 6-well plates (2 × 10^6^/well for MDAMB361, 1.5 × 10^6^/well for other lines) and maintained in at 37°C, 5% CO_2_. After 24 hours, viable cell number was determined by Trypan Blue exclusion, and fresh media was added to replicates placed in a standard incubator (normoxia) and in hypoxic conditions (0.1% O_2_, Whitley H35 Hypoxystation). Viable cells were counted at various times, as indicated.

### Generation of RNF213 mutants

RNF213 mutants were generated by PCR or by using fragments synthesized by Genewiz/IDT. Fragments containing the desired mutation were sequenced, digested, and cloned into wild-type RNF213 plasmids by using T4 DNA ligase or via Gibson assembly^70^.

### NF-κB reporter assays

Cells co-transfected with an NF-κB reporter construct (p2X-NF-κB-BS-Luc-Firefly, a generous gift of Dan Littman, NYU Grossman School of Medicine) and pCMV-Renilla, were subjected to 0.1 % O_2_ for the indicated times. Firefly and Renilla luciferase was measured using the Dual-Luciferase® Reporter Assay System (Promega) according to the manufacturer’s instructions. Reporter activity was quantified as the ratio of Firefly/Renilla signal, and all signals were normalized to the signal in parental BT474 cells.

## IL-1β assays

Cell supernatants were collected at the indicated times, and IL-1β levels were quantified by using the Lumit™ IL-1β Immunoassay (Promega), according to manufacterer’s instructions.

### Biochemical Assays

#### Immunoblotting and immunoprecipations

Cells were lysed in RIPA (total cell lysates, ALFA-tag precipitations) or NP40 buffer (co-immunoprecipitations); buffer compositions and immunoprecipitation conditions have been described^24^ (also see **Method details**). ALFA-tagged proteins were recovered using a single domain camelid antibody bound to magenetic beads (NanoTag Biotechnologies). Antibody details are in Supplementary Table 3.

#### Proximity-based labelling

Cells were cultured in standard media but with 10% tetracycline (Tet)-free FBS (Clonetech) for at least 1 week. Tet-free/biotin-free FBS was generated by incubating Tet-free FBS with sterile streptavidin beads overnight followed by centrifugation to remove the beads. N-Turbo, C-Turbo, and Turbo proteins were induced by adding 1 μg/ml Dox to cells for 36 hours. For the last ten min. of incubation, 500 μM of biotin was added to the medium. Labelling reactions was terminated by removing the medium and adding ice-cold PBS to plates, which were kept on ice. Lysates were collected in RIPA buffer without Na-deoxycholate, and biotin-labelled proteins were collected by incubating with high capacity streptavidin agarose beads for 4 hours at 4°C. Comparable protein amounts (assessed by silver stain and avidin blotting) were analyzed by MS.

### Purification of RNF213 and RNF213 variants

HeLa *RNF213*-KO cells expressing RNF213 or RNF213 variants (except RNF213^H4509A^) were induced with 0.5 μg/ml Dox for 4 days. The 3x-FLAG-RNF213^H4509A^ variant was transiently transfected into HeLa *RNF213*-KO cells, as the stable expression of this variant was low. Lysates from ten 15 cm plates were subjected to anti-FLAG immunoprecipitation. RNF213 species were eluted with 3x-FLAG peptide and concentrated by passage through an Amicon Ultra-4 Centrifugal Filter (100 kDa cut-off).

### In vitro ubiquitylation assays

Ubiquitylation assays were performed as described.^24^ Briefly, 0.25μM of purified RNF213 or RNF213 variant were incubated with 2.5μM GST-SPATA2 (Abnova) or 2.5μM 6xHis-CYLD (R&D) in E3-Ligase buffer (R&D) along with 50μM HA-Ub, 100nM UBE1 (E1), 1μM UBE2L3 (E2), and 10mM Mg-ATP (all from Boston Biochem) for 30 min. in a total reaction volume of 25 μl. Reactions were terminated by adding SDS–PAGE sample buffer followed by incubation for 5 min. at 95°C.

### In vitro kinase assays

Purified full-length 3x-FLAG-tagged RNF213 or RNF213^Y1275F^ were mixed with varying amounts of bacterially produced His-ABL1 (as indicated) and incubated with 600ng of RNF213 in kinase assay buffer (50mM HEPES [pH 7.5], 10mM MgCl_2_, 2.5mM DTT, 75μM ATP) for varying times, as indicated. Final reaction volumes were 20μl. Reactions were terminated by adding SDS–PAGE sample buffer followed by incubation for 5 min. at 95°C.

### Mass spectrometry

Proteins collected on streptavidin beads were digested with sequencing grade trypsin and analyzed by LC/MS/MS using a Thermo Scientific EASY-nLC 1200 and a Thermo Scientific Orbitrap Eclipse Mass Spectrometer MS/MS spectra were searched against the Uniprot human database with the addition of the sequences of RNF213 with TurboID at the N-and C-terminus using Sequest within Proteome Discoverer 1.4. MS/MS spectra were also searched using Byonic by Protein Metrics specifically to identify phosphorylation sites on RNF213. See Extended Methods for details.

### Xenografts and histology

All animal experiments were approved by, and conducted in accordance, with the procedures of the Institutional Animal Care and Use Committee (IACUC) at NYU School Of Medicine (Protocol no. 170602). Six week-old female Balb/c athymic nude mice were implanted with a 60-day release pellet containing 0.72mg 17β-Estradiol (Innovative Research of America). One week later, BT474 cells and BT474 derivatives (1 × 10^6^) or MDAMB361 and MDAMB361 derivatives (2 × 10^6^) resuspended with 50μl of Matrigel (BD), were injected subcutaneously into the right flanks of these mice. Length and width of tumors were measured weekly using Vernier calipers. Tumor volume was calculated as length x width^2^ x 0.52.

For histology and immunohistochemistry, tumors were fixed in 4% paraformaldehyde (PFA) and embedded in paraffin. Sectioning and staining were performed by Perlmutter Cancer Center Experimental Pathology Core.

### Quantification and statistical analysis

Graphs were generated and statistical analyses were performed using GraphPad Prism9. Multiple groups were compared by two-way ANOVA, with additional Tukey multiple comparisons test. Two group comparisons were evaluated by two-tailed t test. Values represent the mean of experiments, with error bars representing S.E.M..

## Extended Methods

### Mass Spectrometry

Proteins bound to streptavidin agarose beads were washed with 100mM NH_4_HCO_3_ (pH 8), reduced with 2.5uL of 0.2M dithiothreitol at 57C for 1 hour, alkylated with 2.5uL of 0.5M IAM for 45 min. at room temperature in the dark, and digested overnight at room temperature with 500ng sequencing grade modified trypsin (Promega). After digestion, samples were acidified to a final concentration of 0.5% trifluoroacetic acid (TFA). A C18 cleanup was performed on all samples using Ultra-Micro SpinColumns (Harvard Apparatus). Briefly, samples were loaded onto equilibrated spin columns, rinsed with 0.1% TFA, and eluted with 40% acetonitrile/0.5% acetic acid and 80% acetonitrile/0.5% acetic acid. Organic solvent was removed using a SpeedVac concentrator, and all samples were reconstituted in 0.5% acetic acid.

One-third of each sample was analyzed individually by LC/MS/MS. Samples were separated online using a Thermo Scientific EASY-nLC 1200, where solvent A was 2% acetonitrile/0.5% acetic acid, and solvent B was 80% acetonitrile/0.5% acetic acid. A 60-min. gradient from 5-35%B was applied to all samples. Peptides were gradient-eluted directly to a Thermo Scientific Orbitrap Eclipse Mass Spectrometer. High resolution, full MS spectra were acquired with a resolution of 240,000, an AGC target of 1e6, a maximum ion time of 50 ms, and a scan range of 400 to 1500 m/z. Following each full MS scan, data-dependent HCD MS/MS spectra collected with low resolution at top speed with a 3ms cycle time. All MS/MS spectra were collected with a rapid scan in the ion trap, an AGC target of 2e4, a maximum ion time of 18 ms, one microscan, 0.7 m/z isolation window, auto scan range mode, and NCE of 27. MS/MS spectra were searched against the Uniprot human database using Sequest within Proteome Discoverer. 1.4. MS/MS spectra were also searched with Byonic by Protein Metrics against a database containing just RNF213 and common contaminants to identify phosphorylation sites. For TurboID analyses, beads were washed two times with 100mM NH_4_HCO (pH 8). Samples were reduced with 2uL of 0.2M dithiothreitol at 57C for 1 hour and alkylated with 2uL of 0.5M IAM for 45 min. at room temperature in the dark. Samples were digested overnight at room temperature with 250ng sequencing trypsin. After digestion, samples were acidified to a final concentration of 0.5% trifluoroacetic acid (TFA), and cleaned up on C18 columns as above. One-third of each sample was analyzed individually by LC/MS/MS. LC conditions were identical to those above. High resolution full MS spectra were acquired with a resolution of 70,000, an AGC target of 1e6, a maximum ion time of 120 ms, and scan range of 400 to 1500 m/z. Following each full MS were 20 data-dependent high resolution HCD MS/MS spectra. All MS/MS spectra were collected with resolution of 17,500, an AGC target of 5e4, a maximum ion time of 120 ms, one microscan, 2 m/z isolation window, fixed first mass of 150 m/z, and NCE of 27. MS/MS spectra were searched against the Uniprot human database with the added sequences of RNF213 with the TurboID on the N-and C-terminus using Sequest within Proteome Discoverer 1.4.

## Extended Data Figure Legends

**Extended Data Figure 1. Y1275 phosphorylation is regulated by PTP1B and controls RNF213 oligomerization.**

(a) PTP1B inhibitor sensitizes multiple breast cancer subtypes to hypoxia. The indicated cell lines were treated with the PTP1B inhibitor claramine, placed in 0.1% O_2_ for 55 hours, and viable cells were counted.

(b) Spectrum supporting phosphorylation of RNF213-Y1275 on peptide EAAEPLSEPKEDQEAAELLSEPEEESERHILELEEVYDYLYQPSYR, from 3x-FLAG-RNF213-TurboID-enriched samples. The HCD MS/MS spectra of sextuply charged m/z 922.9244 ion recorded in the ion trap of an Orbitrap Eclipse is shown. Observed N-terminal sequence ions (b-ions) are shown in blue, and C-terminal fragment ions (y-ions) are shown in red. Backbone cleavages are indicated in the peptide sequence, and the phosphorylated tyrosine is indicated (red).

(c) Y1275 phosphorylation affects RNF213-PTP1B interaction. AU565 *RNF213*-KO-*PTPN1*-KD cells with or without stable expression of doxycycline (Dox)-inducible, HA-*Ptpn1^D/A^* were transiently transfected with 3x-FLAG-*RNF213* or 3x-FLAG-RNF213^Y1275F^ and exposed to 0.5 μg/ml Dox (+) or left untreated (-). Lysates (top) were subjected to anti-HA immunoprecipitation and immunoblotting for the indicated proteins (bottom). ERK2 serves as a loading control.

(d) RNF213**-**Y1275 is also phosphorylated in AU565 cells. AU565 *RNF213*-KO-*PTPN1*-KD cells were transiently transfected with 3x-FLAG-*RNF213* or 3x-FLAG-*RNF213^Y^*^1275^*^F^*. Lysates were subjected to anti-FLAG immunoprecipitation and immunoblotting, as indicated, and pY/FLAG signal (each in arbitrary units) was quantified.

(e) Y1275 phosphorylation promotes RNF213 oligomerization. AU565 *RNF213*-KO-*PTPN1*-KD cells were transiently transfected with expression vectors for *GFP*-*RNF213* and 3x-FLAG-*RNF213* or 3x-FLAG-*RNF213^Y^*^1275^*^F^*. Lysates were subjected to anti-FLAG immunoprecipitation/immunoblotting. Vinculin serves as a loading control.

Error bars represent ± S.E.M.. a, d (lower) are pooled results from three biological replicates; all other data except (b) are representative of these replicates. a, e were evaluated by two-way ANOVA, whereas d was analyzed by ratio paired two-tailed t-test, \*\**P*<0.01, \*\*\**P*<0.001, \*\*\*\**P*<0.0001, ns=non-specific band.

**Extended Data Figure 2. ABL phosphorylates RNF213 on Y1275.**

(a) SKBR3 and SKBR3 *PTPN1*-KO cells were treated with the indicated inhibitors. Cells were subjected to 0.1% O_2_ for 36 hours, and viable cells were counted.

(b) Dasatinib inhibits RNF213-PTP1B interaction. AU565 *RNF213*-KO-*PTPN1*-KD cells expressing substrate trapping mutant HA-*Ptpn1*^D/A^ were transiently transfected with expression vector for 3x-FLAG-*RNF213*, then treated with dasatinib or vehicle (DMSO) for 24 hours. Lysates were subjected to anti-HA immunoprecipitation/immunoblotting.

(c) Dasatinib inhibits tyrosine phosphorylation of RNF213. AU565 *RNF213*-KO-*PTPN1*-KD cells were transiently transfected with 3x-FLAG-*RNF213* followed by treatment with DMSO or dasatinib for 24 hours. Lysates were subjected to anti-FLAG immunoprecipitation/ immunoblotting.

(d) Purity of 3x-FLAG-RNF213 and 3x-FLAG-RNF21^Y1275F^ proteins (see Methods)

(e) ABL phosphorylates RNF213 on Y1275. Purified 3x-FLAG-RNF213 and 3x-FLAG-RNF213^Y1275F^ were phosphorylated *in vitro* with recombinant His-ABL1 for the indicated times (see Methods) (f) ABL1/ABL2 promote RNF213-mediated hypoxia hypersensitivity in *PTPN1*-KO cells. MDAMB361 and MDAMB361 *PTPN1*-KO cells were transfected with siRNAs targeting *ABL1* and/or *ABL2* or control non-targeting siRNA. Cells were subjected to 0.1% O_2_ for 48 hours, and viable cells were counted (top). Immunoblot shows knockdown efficiency (bottom).

Error bars represent ± S.E.M. All panels (a-f) are representative results from two independent experiments, except for the quantifications in a, c, f, which are pooled data from triplicates. a, f were evaluated by two-way ANOVA; C was analyzed by ratio paired two-tailed t-test \*\**P*<0.01, \*\*\*\**P*<0.0001, ns=non-specific band.

**Extended Data Figure 3. Sample preparation for proximity-based labelling screen.**

(a) HeLa FlpIn-TRex cells reconstituted with Control Turbo, N-turbo, or C-turbo were induced with 0.5 μg/ml of Dox. Post-biotin labelling (see Methods), lysates (left) were subjected to streptavidin affinity purification (right) and immunoblotting.

(b) Silver-stained SDS-PAGE gel showing that comparable amounts of streptavidin-purified proteins were submitted for mass spectrometry.

(c) One-sided volcano plot showing fold-change and SAINT score of proteins identified in N-Turbo versus Control Turbo. Proteins with SAINT Score >5% FDR are highlighted.

(d, e) GO pathway analysis showing top 10 enriched molecular functions of C-Turbo (d)-and N-Turbo (e)-interacting proteins.

**Extended Data Figure 4. CYLD/SPATA2 are relevant substrates of RNF213 in mediating hypoxia hypersensitivity.**

(a, b) SKBR3 and SKBR3 *PTPN1*-KO cells, with or without CYLD (a) or SPATA2 (b), were subjected to 0.1% O_2_ for the indicated times, and viable cells were counted (top). Immunoblots (bottom) show absence of CYLD/SPATA2 in stable pools generated with the indicated sgRNAs.

(c) RNF213 promotes SPATA2 and CYLD ubiquitylation. AU565 and AU565 *RNF213*-KO cells were co-transfected with expression contructs for HA-UB and ALFA-*SPATA2* or ALFA-*CYLD* and treated with claramine or vehicle (-), as indicated. Lysates (left) were subjected to anti-ALFA immunoprecipitation/immunoblotting (right).

(d) Purity of 3x-FLAG-RNF213, 3x-FLAG-RNF21^H4509A^, 3x-FLAG-RNF21^H4014A^, 3x-FLAG-RNF21^H4014A/H4509A^ after indicated steps (see Methods)

(e, f) Effects of RNF213 mutants on global ubiquitylation. Hela FlpIn-TRex *RNF213*-KO (e) and BT474 *RNF213*-KO (f) cells were co-transfected with expression contructs for HA-UB and 3x-FLAG-tagged *RNF213* or -*RNF213^H^*^4014^*^A^* --*RNF213^H^*^4509^*^A^* or -*RNF213^H^*^4014^*^A/H^*^4509^*^A^*. Cells were subjected to 0.1% O_2_ for the indicated times, and lysates were subjected to immunoblotting. Error bars represent ± S.E.M.. a and b are pooled results from three biological replicates; data in c, e and f are representative of replicates, \*\*\*\**P*<0.0001, two-way ANOVA, ns=non-specific band.

**Extended Data Figure 5. RNF213 RING mutants/highly penetrant MMD SNPs activate RZ domain and require ATPase activity.**

(a-c) AU565 *RNF213*-KO cell reconstituted with RNF213^WT^ or the indicated RNF213 mutants, parental AU565 cells, and AU565 *RNF213*-KO cells with or without *PTPN1* deletion, were subjected to 0.1% O_2_ for 40 hours. Immunoblotting was performed for the indicated proteins. Each blot is representative of two biological replicates, ns=non-specific band.

**Extended Data Figure 6. Highly penetrant MMD SNPs activate RZ domain.**

(a) BT474 *RNF213*-KO cell reconstituted with RNF213^WT^, the indicated RNF213 mutants, parental BT474 cells, and BT474 *RNF213*-KO cells with or without PTP1B, were subjected to 0.1% O_2_ for 40 hours. Immunoblotting was performed for the indicated proteins.

(b) AU565 *RNF213*-KO cell reconstituted with RNF213^WT^ or the indicated RNF213 mutants, parental AU565 cells, and AU565 *RNF213*-KO cells with or without *PTPN1* deletion, were subjected to 0.1% O_2_ for 40 hours. Immunoblotting was performed for the indicated proteins.

(c) RZ and ATPase domains of RNF213 are required for hypoxia hypersensitivity of AU565 *PTPN1*-KO cells. Cells prepared as in (b) and Extended Data Fig. 5 a-c were subjected to 0.1% O_2_, and viable cells were counted after 48 hours.

Error bars represent ± S.E.M.. (n = 3 biological replicates, each panel); \*\*\**P*<0.001, \*\*\*\**P*<0.0001, two-way ANOVA

**Extended Data Figure 7. *PTPN1*-KO cells die via RNF213-and LUBAC-dependent pyroptosis.** (a) Wild-type, but not RING-dead, RNF213 requires LUBAC for CYLD/SPATA2 degradation. AU565 *RNF213*-KO cells reconstituted with wild-type *RNF213* or *RNF213^H^*^4014^*^A^*, parental AU565 cells. and AU565 *RNF213*-KO cells with or without *PTPN1* deletion were transfected with *RBCK1* or control siRNAs. Cells were subjected to 0.1% O_2_ for 40 hours, lysed, and immunoblotted as indicated (top). *RBCK1* knockdown was confirmed by RT-qPCR (bottom).

(b) RING-dead RNF213 bypasses LUBAC requirement. The indicated cells were subjected to 0.1 % O_2_ for 48 hours, and viable cell numbers were counted.

(c) RNF213 promotes NF-κB activity. The cells described in Extended Data Fig. 6c were transfected with an NF-κB reporter construct (see Methods) and subjected to 0.1% O_2_ for 40 hours. Firefly and Renilla luciferase signals were measured, reporter activity was quantified as the ratio of Firefly/Renilla signal, and all signals were normalized that from parental AU565 cells.

(d, e) Hypoxic cell death in *PTPN1*-KO cells is caspase-dependent. SKBR3 and SKBR3 *PTPN1*-KO (d) and MDAMB361 and MDAMB361 *PTPN1*-KO (e) cells were treated with the indicated inhibitors, placed in 0.1% O_2_ for 40 (d) or 48 (e) hours, and cell viability was assessed.

(f, g) Hypoxic cell death in MDAMB361 *PTPN1*-KO cells requires inflammatory caspases and the inflammasome. As in (e), except with/without VX765 (f) or MCC950 (g).

(h) GSDMD promotes hypoxic cell death of *PTPN1*-KO cells. Pools of MDAMB361 and MDAMB361 *PTPN1*-KO cells, with or without *GSDMD* deletion, were subjected to 0.1% O_2_, and cell viability was measured at 48 hours (top). Immunoblot indicates efficiency of *GSDMD* knockout (bottom).

(i) Hypoxia-hypersensitive cells release IL-1β. Cells prepared as in Extended Data Fig. 6c were subjected to 0.1% O_2_ for 48 hours, and IL-1β in supernatants was analyzed by immunoassay.

In all panels, error bars represent ± S.E.M. (n = 3 biological replicates), and each blot is representative of two biological replicates. \*\*\**P*<0.001, \*\*\*\**P*<0.0001, two-way ANOVA, ns=non-specific band.

**Extended Data Figure 8. Hypoxia-evoked ER stress activates the inflammasome for RNF213-triggered pyroptosis.**

(a) Hypoxia hypersensitivity is dependent on O_2_ levels. BT474 and BT474 *PTPN1*-KO cells were subjected to varying levels of O_2_ (0.1, 0.2, 0.5 and 1% O_2_) for 55 hours, and viability was assessed.

(b) Hypoxia hypersensitivity is dependent on O_2_ level. MDAMB361 and MDAMB361 *PTPN1*-KO cells were subjected to varying levels of O_2_ (0.1, 0.2, 0.5 and 1% O_2_) for 55 hours, and viability was assessed.

(c, d) ER stress promotes hypoxia hypersensitivity. MDAMB361 and MDAMB361 *PTPN1*-KO cells treated with the IRE1α inhibitor 4μ8C (B) or the PERK inhibitor GSK2606414 (C), were subjected to 0.1 % O_2_ for 48 hours, and viability was assessed.

(e) ER stress causes pyroptosis in CYLD/SPATA2-deficient cells in normoxia. MDAMB361 cells, with or without CYLD/SPATA2, were treated with tunicamycin (10 μM) or thapsigargin (5 μM) for 40 hours and viability was assessed. Error bars represent ± S.E.M.. (n = 3 biological replicates), \*\*\*\**P*<0.0001, two-way ANOVA.

**Extended Data Figure 9. The PTP1B/RNF213/CYLD/SPATA2 pathway controls HER2+ breast cancer tumorigenesis**

(a, b) MDAMB361 and MDAMB361 *PTPN1*-KO cells, with or without *CYLD*-KO, were injected subcutaneously into athymic *nu/nu* mice (6/group). Tumor size was measured weekly for 6 weeks (a). Residual tumors were excised, and fixed and stained with H&E (b). * in (b) indicates representative necrotic regions.

(c) Left panel: ABL1/2-catalyzed phosphorylation of RNF213 at Y1275 promotes RNF213 oligomerization; PTP1B dephosphorylates this site. Oligomerized RNF213, via its RZ domain, ubiquitylates CYLD and SPATA2, and further ubiquitylation, catalyzed by the LUBAC complex, promotes CYLD/SPATA2 degradation.

Left middle panel: ATPase-dead RNF213 cannot oligomerize and does not promote hypoxia hypersensitivity.

Right middle panel:RING-dead RNF213 enhances RZ-mediated ubiquitylation of CYLD/SPATA2 and leads to CYLD and SPATA2 degradation even in the absence of LUBAC. Degradation of CYLD/ SPATA2 leads to increased NF-κB signaling, priming pyroptosis. Hypoxia-associated ER-stress provides the “second signal” to activate the inflammasome formation, leading to RNF213-driven pyroptotic cell death.

Right panel: RZ-dead or RING/RZ-dead RNF213 cannot ubiquitylate CYLD/SPATA2 and does not promote hypoxia hypersensitivity in response to PTP1B inhibition.

