## Supplementary figures and images for "Identification of a Novel Hypoxia-induced Inflammatory Cell Death Pathway"

### Extended Data Fig. 1

Extended Data Figure 1

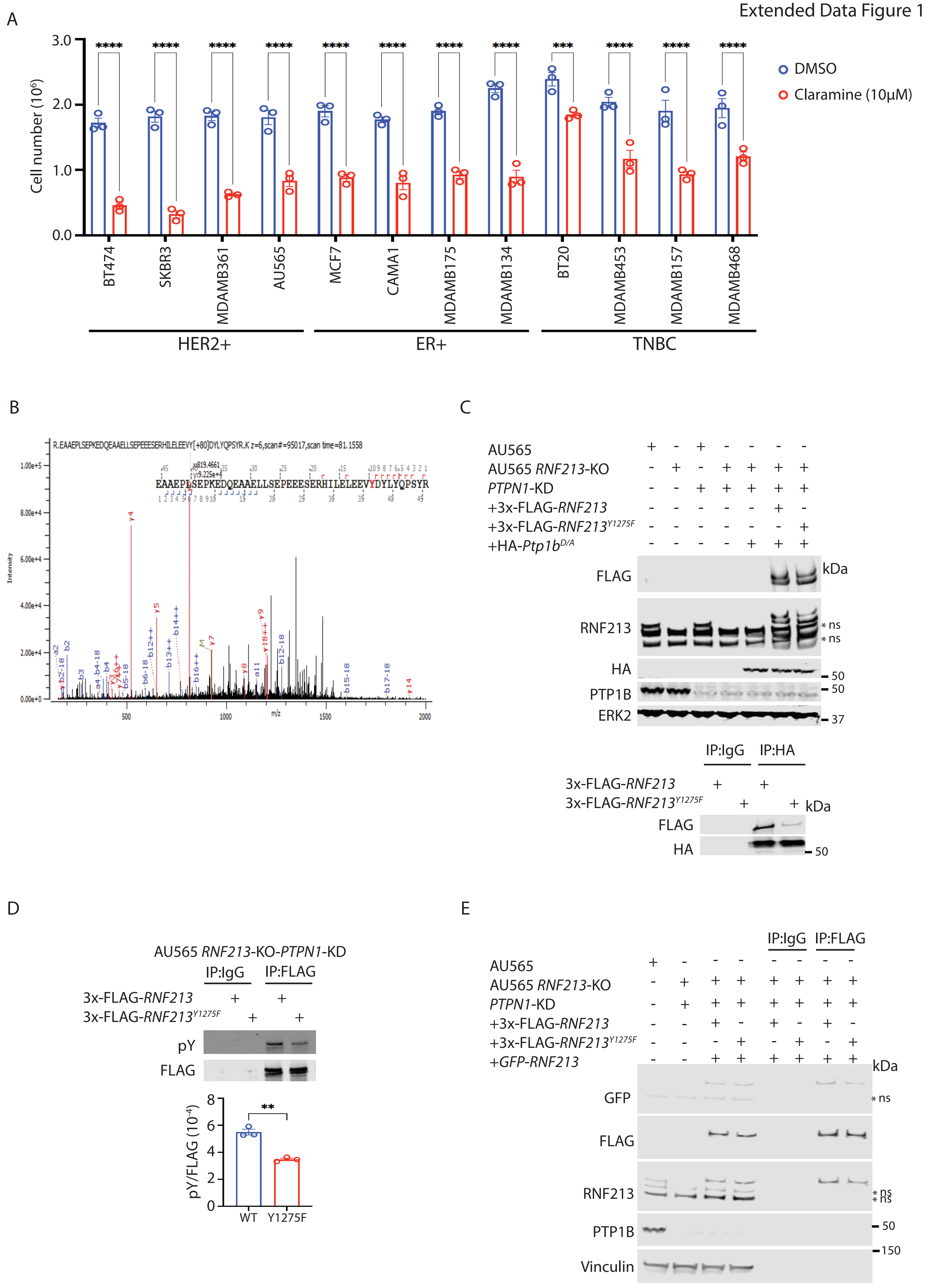

### Extended Data Fig. 2

Extended Data Figure 2

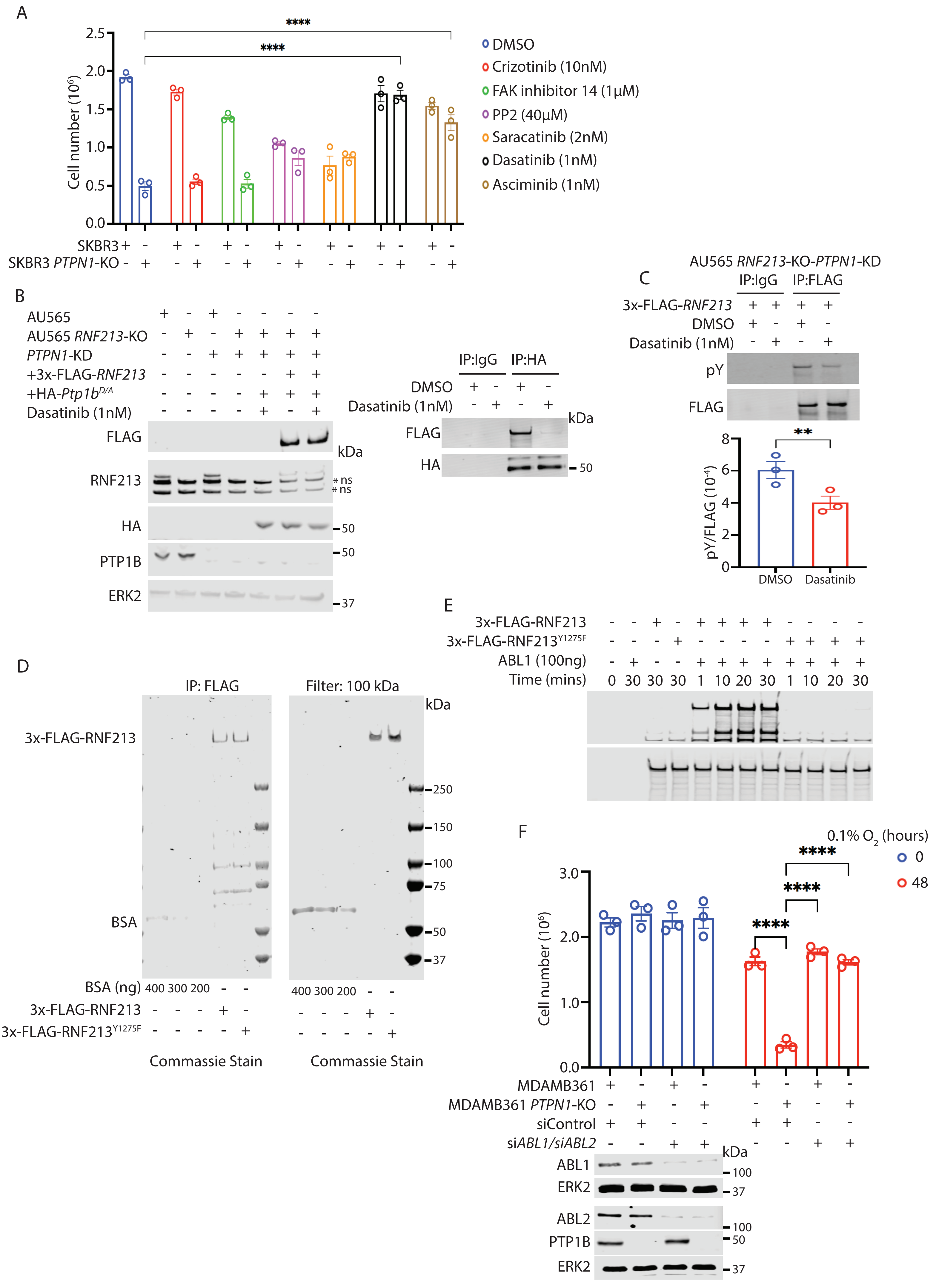

### Extended Data Fig. 3

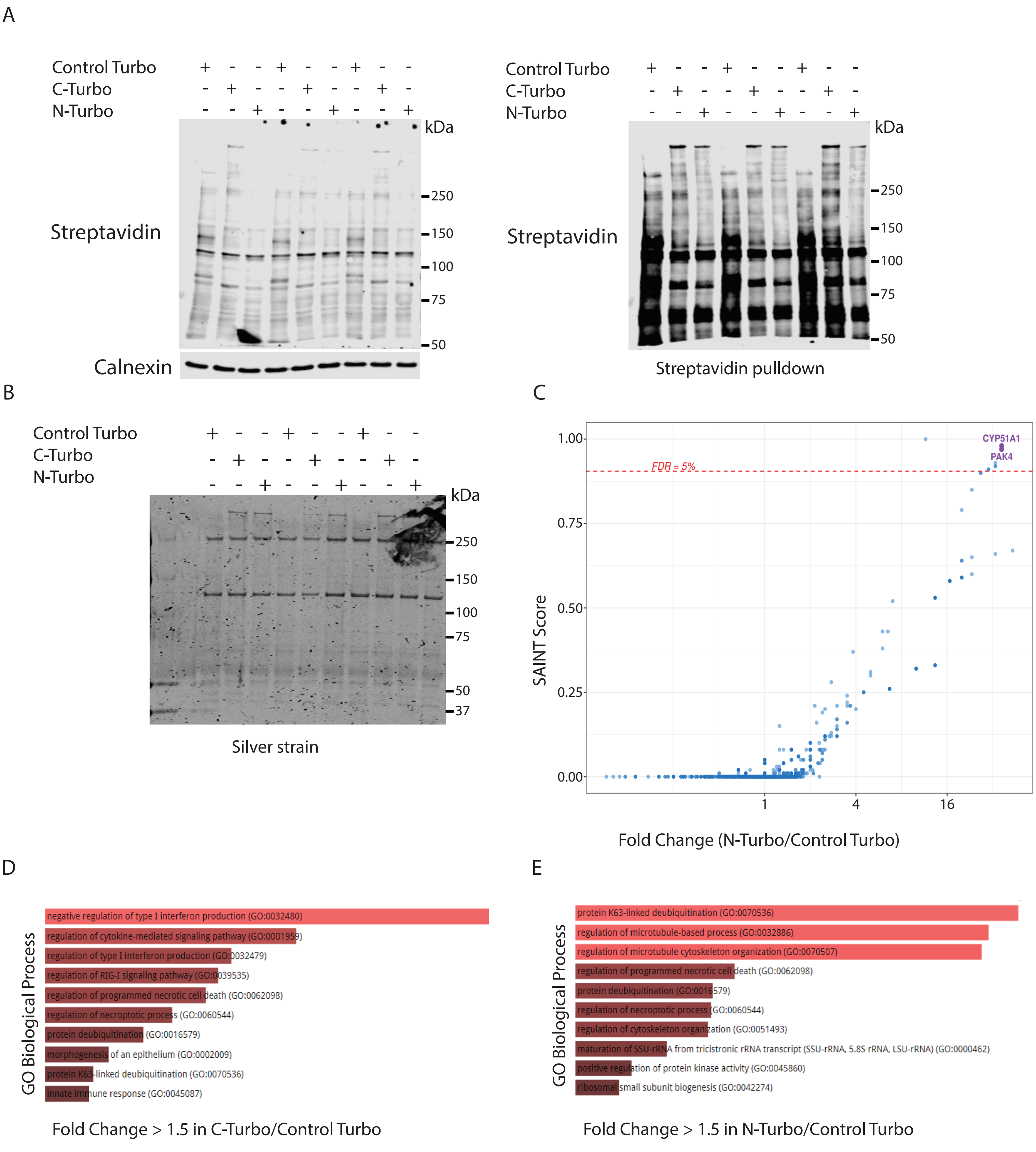

### Extended Data Fig. 4

Extended Data Figure 4

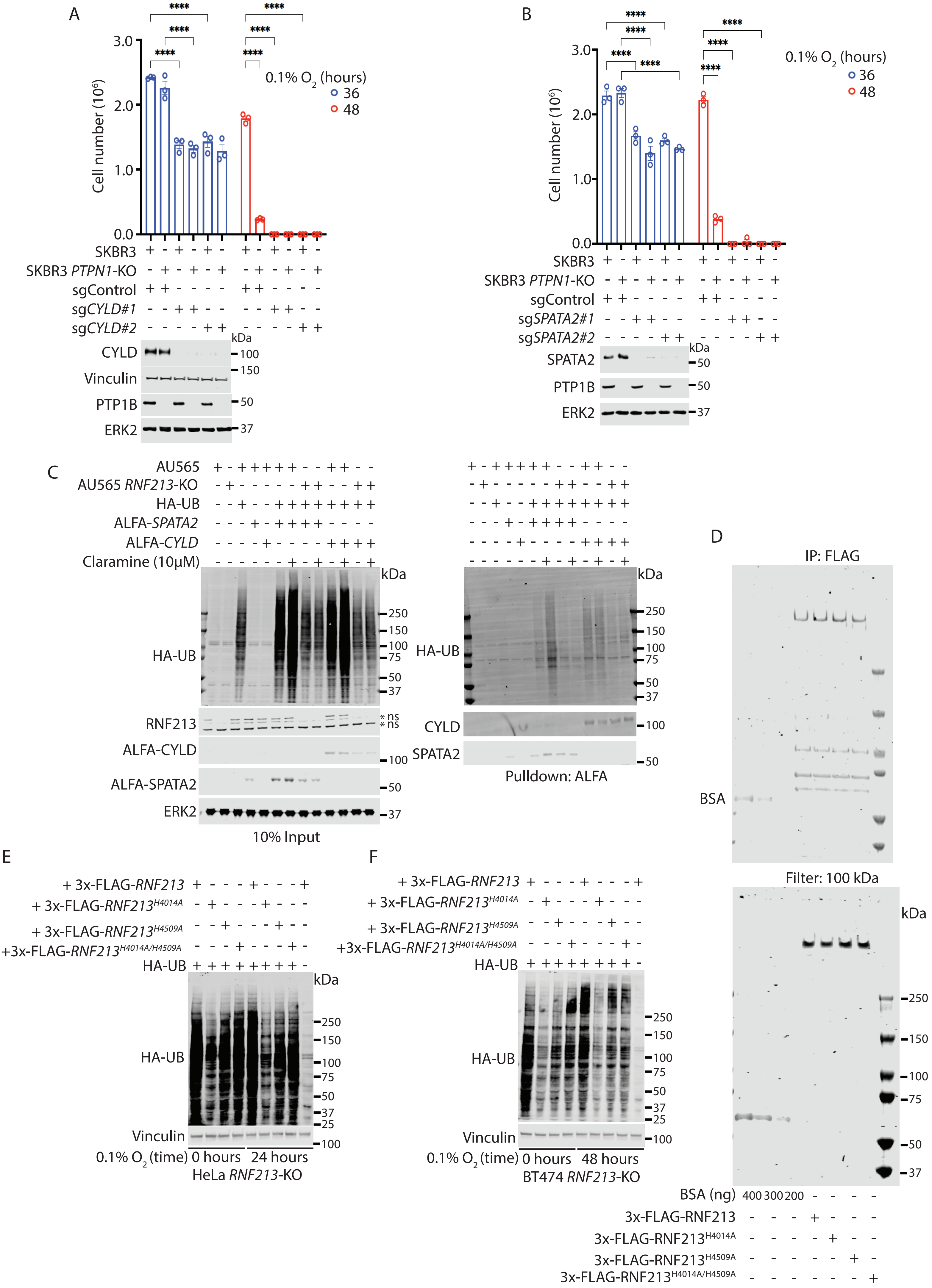

### Extended Data Fig. 5

#### Extended Data Figure 5

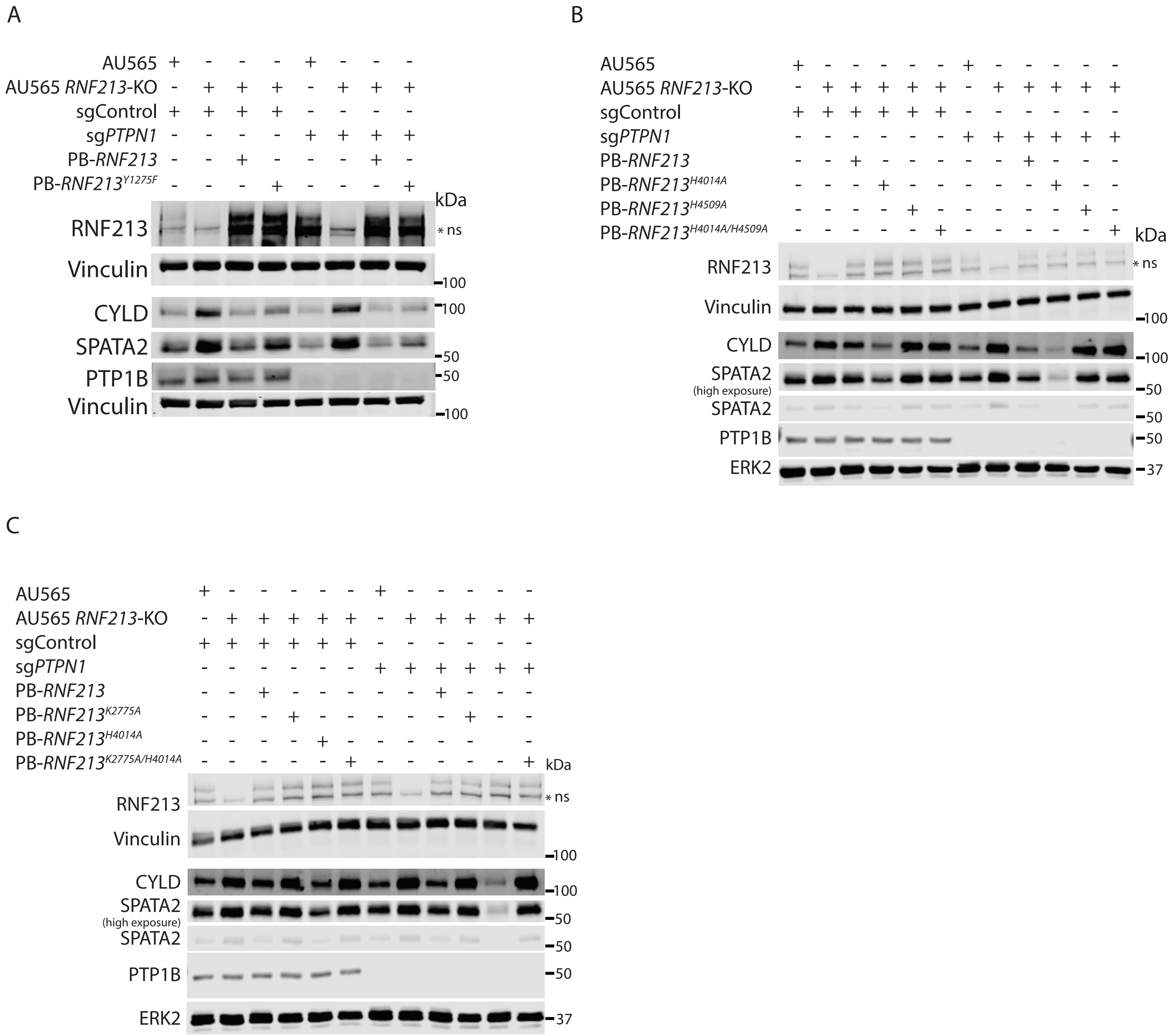

### Extended Data Fig. 6

#### Extended Data Figure 6

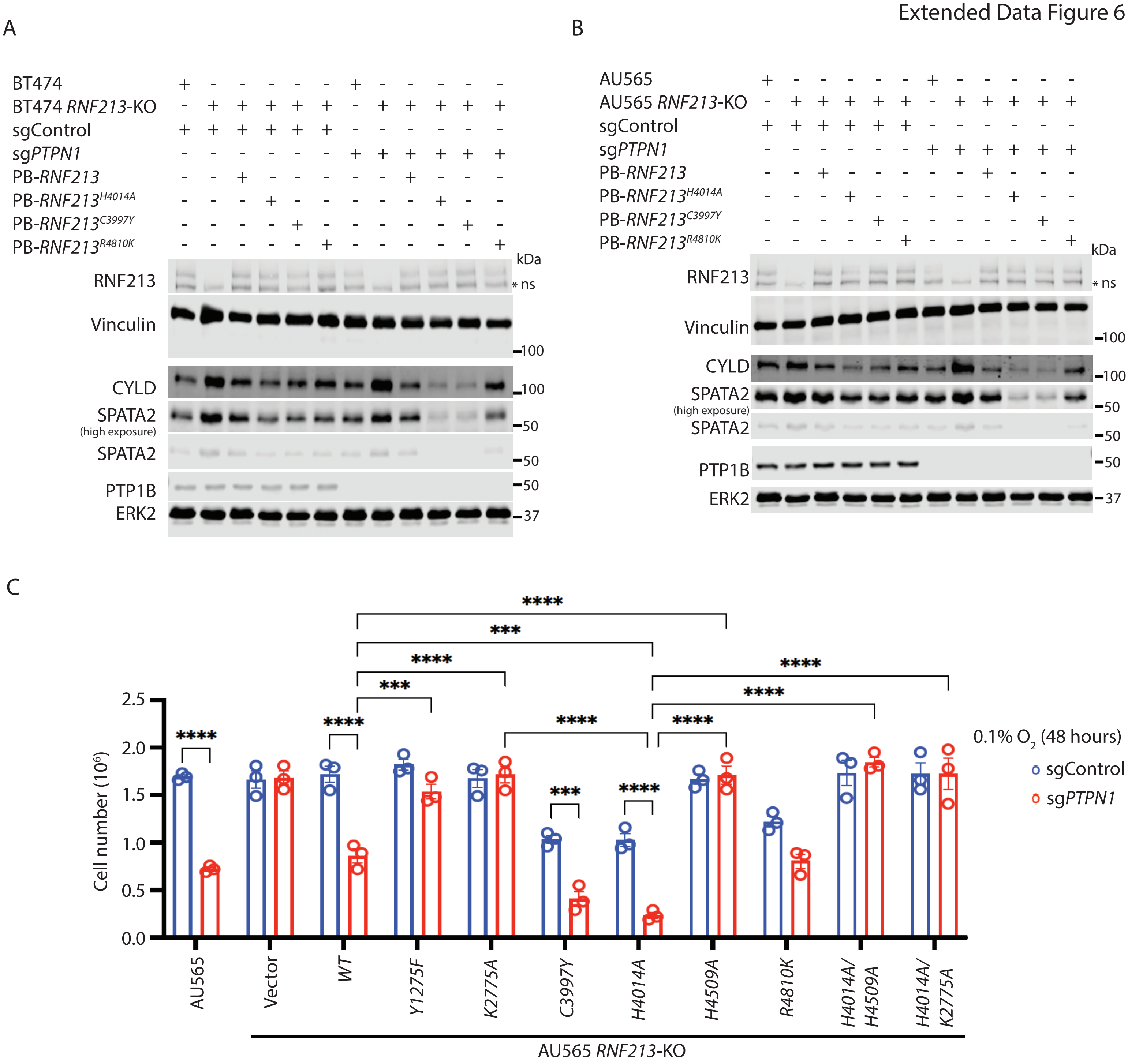

### Extended Data Fig. 7

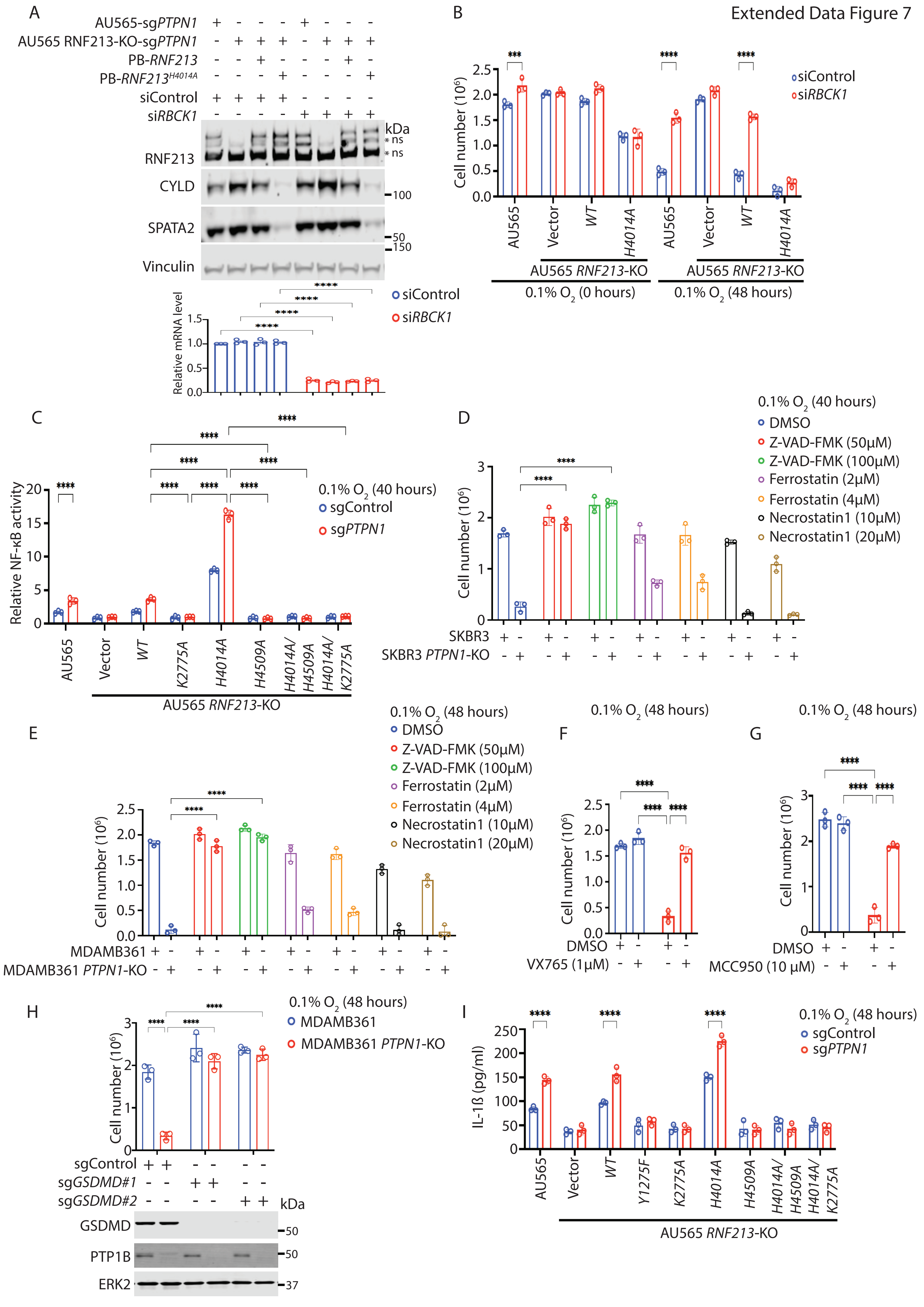

### Extended Data Fig. 8

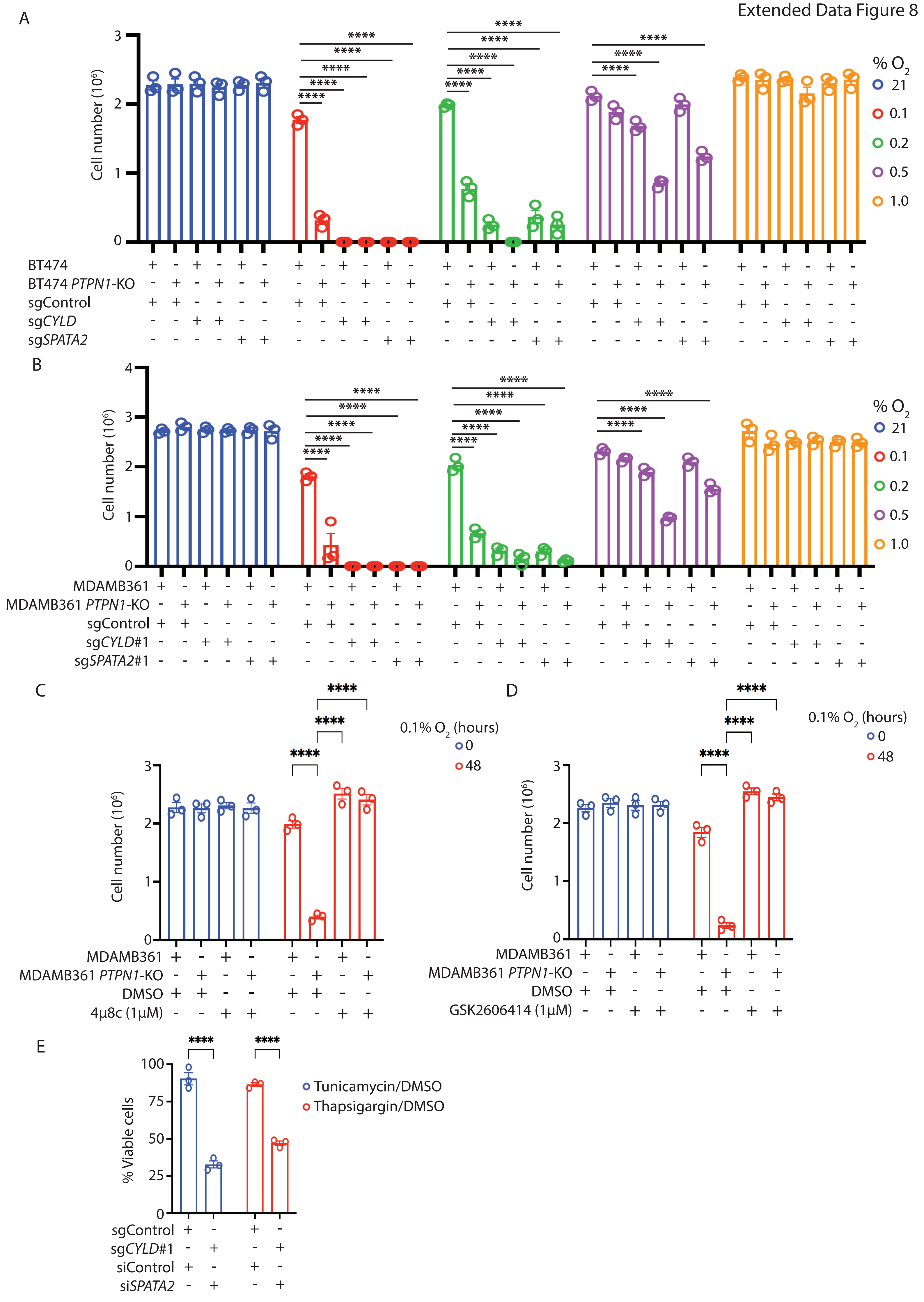

### Extended Data Fig. 9

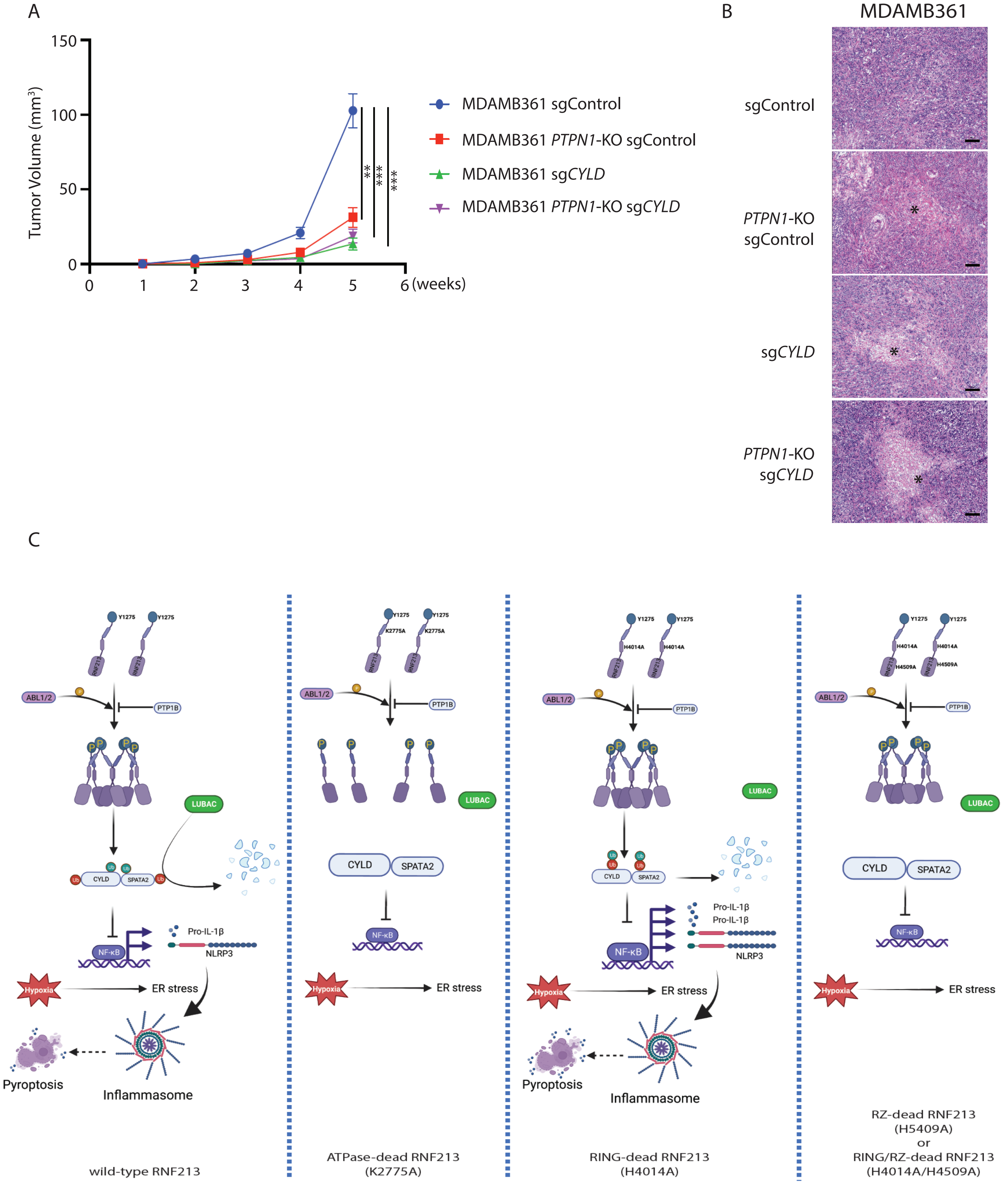
